# Daily glucocorticoids promote glioblastoma growth and circadian synchrony to the host

**DOI:** 10.1101/2024.05.03.592418

**Authors:** Maria F. Gonzalez-Aponte, Anna R. Damato, Tatiana Simon, Nigina Aripova, Fabrizio Darby, Joshua B. Rubin, Erik D. Herzog

## Abstract

Glioblastoma (GBM) is the most common primary brain tumor in adults with a poor prognosis despite aggressive therapy. A recent, retrospective clinical study found that administering Temozolomide in the morning increased patient overall survival by 6 months compared to evening. Here, we tested the hypothesis that daily host signaling regulates tumor growth and synchronizes circadian rhythms in GBM. We found daily Dexamethasone promoted or suppressed GBM growth depending on time of day of administration and on the clock gene, *Bmal1*. Blocking circadian signals, like VIP or glucocorticoids, dramatically slowed GBM growth and disease progression. Finally, mouse and human GBM models have intrinsic circadian rhythms in clock gene expression *in vitro* and *in vivo* that entrain to the host through glucocorticoid signaling, regardless of tumor type or host immune status. We conclude that GBM entrains to the circadian circuit of the brain, which modulates its growth through clock-controlled cues, like glucocorticoids.

## Introduction

Glioblastoma (GBM) is the most common and deadly brain tumor in adults^1,2^. Despite an aggressive treatment paradigm that includes maximal safe surgical resection, radiation plus concomitant and adjuvant chemotherapy with Temozolomide (TMZ), and tumor-treating fields, median survival time post-treatment is 15 months, and 5-year survival is less than 5% after diagnosis^1,3,4^. These devastating trends emphasize the importance of identifying novel therapeutic targets and approaches that can improve outcomes for GBM patients. One recent approach shown to maximize response to chemotherapy and extend survival in cellular and animal models of GBM, as well as in human patients, is administering TMZ in accordance with time of day^5–8^. A recent retrospective clinical study found that taking TMZ in the morning compared to the evening was associated with a 6-month increase in median survival^6^. We recently demonstrated that circadian regulation in expression of the DNA repair enzyme, MGMT, underlies daily rhythms in TMZ sensitivity in cellular and animal models of GBM^8^. These findings suggest that circadian rhythms in GBM may regulate tumor biology and response to therapies.

In addition to therapies that aim to reduce tumor growth, GBM patients are currently treated with corticosteroids around the time of surgery and radiation, and often at the time of terminal progression, to reduce GBM-induced cerebral edema^9–11^. Dexamethasone (DEX) is the drug of choice among synthetic glucocorticoids (GCs, cortisol in humans and corticosterone in rodents) due to its ability to decrease the permeability of the blood brain barrier^9,12–14^, high specificity for the glucocorticoid receptor (GR), long half-life, and high potency^10^. However, DEX has been variably implicated in tumor progression^11^. Several studies have shown tumor suppressive effects of DEX in patients and in various GBM models *in vitro* and *in vivo*^15–19^. Others have demonstrated DEX promotes GBM cell proliferation and a glioma stem cell-like phenotype, decreases host survival, and induces resistance to chemotherapy with TMZ^11,20–25^. None of these studies have controlled for the possibility of glucocorticoid action varying with time of day. Because glucocorticoid release increases several folds each day prior to waking^26–30^, time of DEX administration and daily glucocorticoid secretion may differentially impact GBM progression and outcomes. Here, we hypothesized that DEX and daily glucocorticoid secretion regulates tumor progression dependent on circadian time in GBM.

Unlike other cancers where circadian rhythms tend to be disrupted, well-studied murine, human, and primary GBM models have reliable circadian rhythms in clock gene expression and sensitivity to therapies^5,8,31,32^. This leads to the hypothesis that GBM tumors act as peripheral circadian pacemakers that synchronize (entrain) their daily rhythms to the host to regulate tumorigenic processes. To maintain synchronized circadian rhythms in physiology and behavior, all vertebrates depend on a central circadian pacemaker in the suprachiasmatic nucleus (SCN) of the brain that entrains its daily rhythms in clock gene expression through light and neuropeptides, such as pituitary adenylate cyclase-activating peptide (PACAP) and vasoactive intestinal peptide (VIP)^33–39^. The SCN, in turn, regulates daily rhythms in the rest of the body through signals, including body temperature^40,41^, glucocorticoids^42,43^, and insulin/insulin-like growth factor (IGF) in combination with glucose^44–46^. Among these, glucocorticoids are one of the highest amplitude circadian outputs, and one of the most potent synchronizing cues for circadian clocks in tissues including brain, liver, kidney, and heart^42^. Here we test the hypothesis that daily glucocorticoid signaling synchronizes circadian rhythms in GBM to the host. Altogether, we find that blocking the daily peak of glucocorticoid signaling desynchronizes circadian rhythms in GBM from the host and dramatically slows disease progression in tumor-bearing mice.

## Results

### Dexamethasone promotes GBM growth dependent on circadian time of treatment

DEX is reported to both promote and suppress GBM tumor^11^. As DEX is commonly used to control brain edema in GBM, it is important to clarify what distinguishes growth-promoting from growth suppressive effects. We hypothesized that known circadian variation in daily glucocorticoid secretion and action might underlie the variability in DEX effects. We used the well-characterized human LN229 and murine GL261 GBM cell lines, which have been previously found to have reliable circadian rhythms in clock gene expression and response to TMZ chemotherapy^8^. We transduced LN229 and GL261 GBM cultures with luciferase reporters driven by the promoters of the clock genes *Bmal1* or *Per2* (Figure 1A). Real-time *in vitro* bioluminescence recordings showed that GBM cells have intrinsic daily rhythms in *Bmal1* (B1L, green) and *Per2* (P2L, yellow) expression (Figures 1B, D and S1A-C), with *Bmal1* consistently peaking approximately 10 h before peak *Per2* expression, and a circadian period ranging from 23h to 31h (Figures S1A-B). We next treated LN229-B1L and GL261-B1L cultures with 100nM DEX or vehicle (0.001% ethanol) at either the daily peak or trough of *Bmal1* expression. We found that DEX promoted a 3-fold increase in GBM growth when delivered around peak of *Bmal1*, but suppressed growth by 3-fold at the trough, in both cell lines (Figure 1C, E). To test whether the growth-promoting effects of DEX depend on an intact circadian clock, we used two different viral-mediated short-hairpin RNA constructs to reduce *Bmal1* levels (Figure S2A). We found both knockdown (KD) constructs disrupted the intrinsic circadian clock, as evidenced by decreased amplitude of circadian *Per2* expression in LN229-P2L cells (Figure 1F and S2B-C). We next treated LN229-*Bmal1* KD cells with 100nM DEX or vehicle at two circadian times that corresponded to either the peak or trough of *Bmal1* expression in LN229-B1L cells. We found that *Bmal1* KD abolished DEX-induced tumor growth or suppression (Figure 1G and S2D). We conclude that the circadian clock in GBM cells regulates DEX-induced growth around the daily *Bmal1* peak and suppression around the daily *Per2* peak.

**Figure 1:**
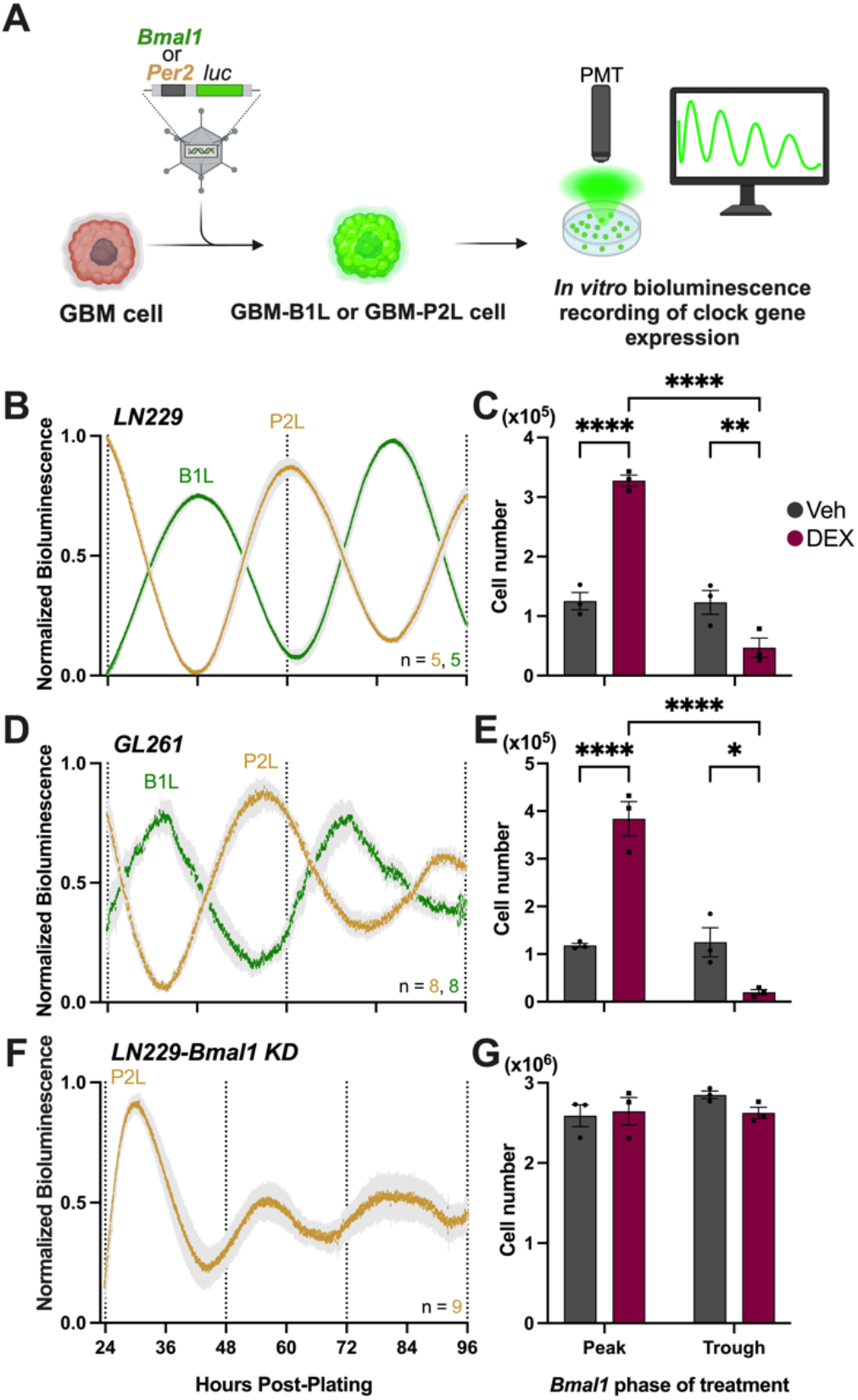
Dexamethasone regulates GBM growth dependent on circadian time of treatment *in vitro* (A) Schematic of cell transduction with luciferase reporters. (B, D) Human LN229 and murine GL261 GBM cell lines transduced with a *Per2-* or *Bmal1-driven* luciferase reporter (P2L and B1L, respectively), show circadian rhythms in clock gene expression *in vitro* (mean±SEM, all recordings had cosine fits with correlation coefficients, CC > 0.9). See also Figure S1. (C, E) Acute treatment with 100nM Dexamethasone at either the peak or trough of *Bmal1* expression *in vitro* shows a 3-fold increase in cell growth when treated at the peak, and a 3-fold growth suppression at the trough (n=3 per condition, mean±SEM, n = 3 per group, two-way ANOVA with Šídák’s multiple comparisons test, ****p < 0.0001, ns p > 0.05). (F) *Bmal1* KD disrupts the circadian clock, as evidenced by reduced amplitude of *Per2* expression in LN229-P2L cells *in vitro* after 48 hours of recording (mean±SEM, all recordings had cosine fits with correlation coefficients, CC = 0.7). (G) LN229-*Bmal1* KD cells acutely treated with 100nM Dexamethasone show no differences in cell growth when treated at either the peak or trough of *Bmal1* expression, relative to LN229-B1L cells, *in vitro* (mean±SEM, n = 3 per group, two-way ANOVA with Šídák’s multiple comparisons test, ns p > 0.05). See also Figure S2.

To evaluate *in vivo* DEX sensitivity as a function of time of day, we implanted either human LN229 cells into immunocompromised nude mice or murine GL261 cells into immunocompetent C57Bl/6NJ mice housed in a light/dark cycle (LD, where lights on is defined as Zeitgeber Time (ZT) 0 and lights off as ZT12). GBM cells expressed either the *Bmal1*- or *Per2*-luciferase reporter. Approximately 10 days after implantation into the basal ganglia, we imaged *in vivo* bioluminescence every 4 h over 36 h from anesthetized mice (Figure 2A). We found reliable daily rhythms in tumor clock gene expression, with *Bmal1* peaking during the day (ZT4) and *Per2* peaking at night (ZT18; Figure 2B-E). Additional recordings of *Bmal1* and *Per2* expression from GL261 or LN229 xenografts in freely moving mice also showed anti-phase circadian rhythms in clock gene expression over 48 h in constant darkness (Figure S3A-F).

**Figure 2:**
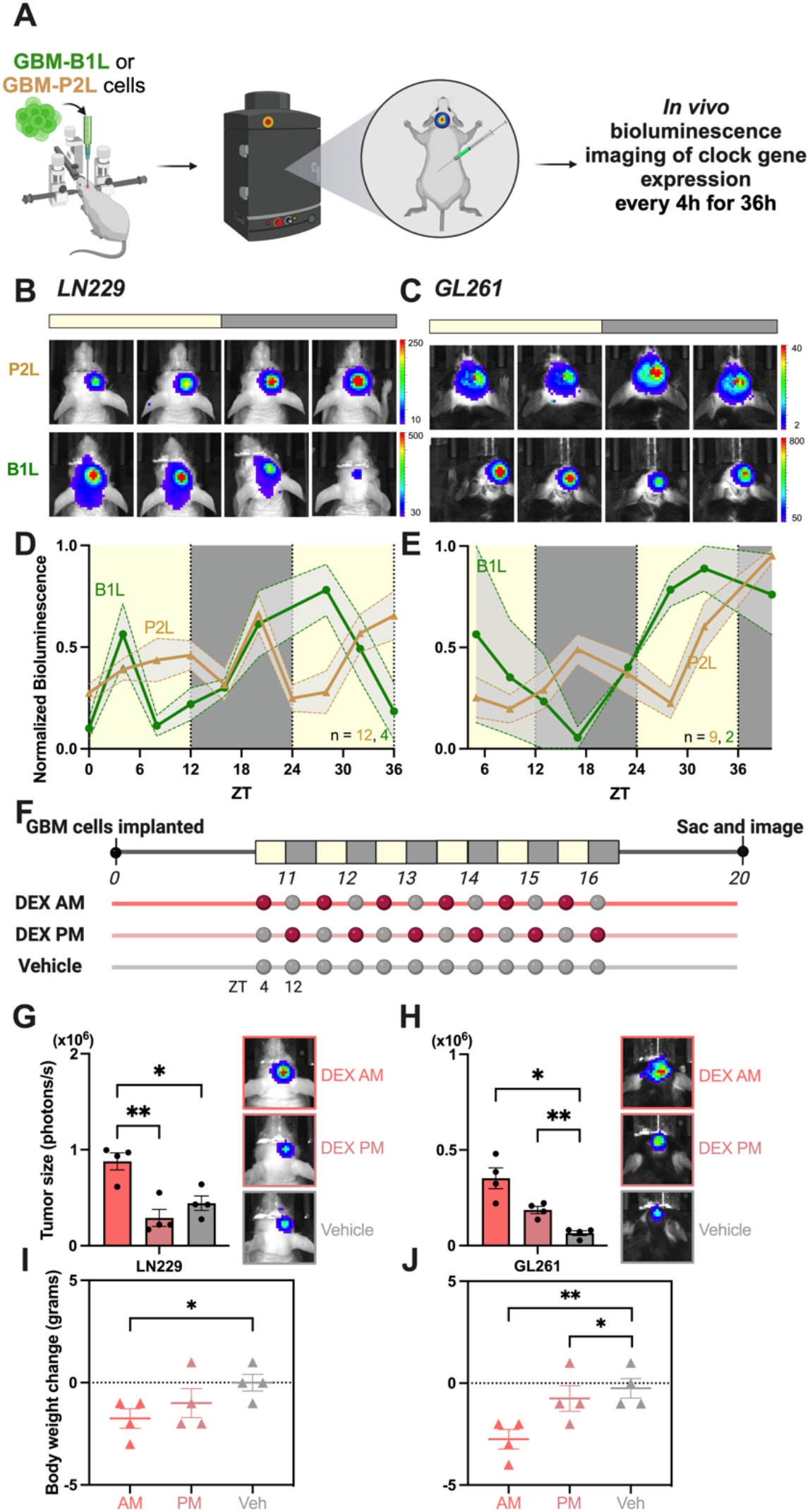
Dexamethasone promotes GBM growth when administered in the morning and at the daily peak of *Bmal1* expression *in vivo* (A) Schematic of orthotopic xenograft into mouse basal ganglia and bioluminescence imaging. (B-C) Representative bioluminescence images of tumor xenografts in mice during the day (ZT0-12) and night (ZT12-24) show high *Bmal1* expression during the day, and high *Per2* expression during the night (BLI counts are x10^3^). (D-E) GBM xenografts show reliable peak *Per2* expression at night and *Bmal1* during the day when implanted in nude or C57BL/6NJ male or female mice (mean±SEM, average traces scored circadian by JTK cycle p < 0.05). See also Figure S3, S4, and S5. (F) Schematic of Dexamethasone treatment paradigm after tumor implantation. (G-H) Dexamethasone treatment promotes LN229 and GL261 tumor growth *in vivo* in a time-dependent manner. A 2- to 5-fold increase in tumor size is observed when treating in the morning (ZT4), compared to evening (ZT12) and vehicle treatments (mean±SEM, n = 4 per group, one-way ANOVA with Tukey’s multiple comparisons test, **p < 0.01, ****p < 0.0001). See also Figure S6. (I-J) Mice bearing LN229 or GL261 tumors treated with Dexamethasone in the morning lost more weight from start to end of the experiment, compared to mice treated in the evening or with vehicle (mean±SEM, n = 4 per group, one-way ANOVA with Tukey’s multiple comparisons test, *p < 0.05, **p < 0.01).

We next tested whether circadian rhythms differ among GBM models or with disease progression *in vivo*. We found similar anti-phase daily rhythms in *Bmal1* and *Per2* expression recorded *in vitro* and from xenografts of a murine model of GBM (NF1^-/-^ DNp53) and a primary GBM isolate (B165) (Figure S4A-H). Consistent with our previous results, *Bmal1* and *Per2* expression peaked during the light and dark phases, respectively, in NF1^-/-^ DNp53 and B165 GBM tumors *in vivo* (Figure S4G-H). Further, imaging of later stage GL261-P2L xenografts showed that tumors grew significantly, and mice lost significant body weight from 11 to 18-days post-implant (Figure S5A). Regardless of the stage of tumor progression, we found that *Per2* still reliably peaked during the night (ZT16; Figure S5B-E).

To assess DEX effects *in vivo* at different times of day, we implanted mice with either human LN229-B1L or murine GL261-B1L cells, and tracked tumor size by *in vivo* bioluminescence imaging. Once tumor growth was established at 11 days post-implant, we delivered 0.5 mg/kg DEX or vehicle (water) by oral gavage for 6 consecutive days at either 4-h (ZT4) or 12-h (ZT12) after daily light onset (corresponding to the peak or trough of tumor *Bmal1* expression, respectively). To control for the effects of mouse handling, we treated all mice with vehicle by oral gavage at the times when they did not receive DEX (Figure 2F). We found that LN229 and GL261 tumor size significantly increased by 5- and 2-fold, respectively, in mice receiving DEX in the morning (Figure 2G-H). Mice treated with DEX in the morning lost significantly more weight than those treated in the evening or with vehicle, indicating greater disease progression (Figure 2I-J). As an additional indicator of tumor proliferation, we collected brain tissue at 20 days post-implant and stained for the proliferation marker Ki67. We found significantly higher Ki67 expression and tumor area in brain sections from mice receiving DEX in the morning, compared to evening or vehicle (Figure S6A-B). Together, these data demonstrate that diverse GBM tumors synchronize their daily cycles of clock gene expression to their host, regardless of host immune status and over the course of disease progression, and that DEX in the morning *in vivo* (around peak *Bmal1* expression) promotes GBM growth and accelerates disease progression.

### Daily glucocorticoid signaling to GBM promotes tumor growth and accelerates disease progression

To test the hypothesis that daily endogenous glucocorticoid signaling regulates GBM progression, we reduced expression of the glucocorticoid receptor (GR) in LN229-P2L and GL261-P2L GBM cells using viral-mediated knock-down (hereafter, GR KD; Figure S7A-B, E-F). GR KD of 2- to 5-fold did not affect intrinsic cell growth or daily *Per2* expression *in vitro* (Figure S7C-D, G-H), but eliminated glucocorticoid-induced cell growth (Figure 3A, F), suggesting that glucocorticoids must act through glucocorticoid receptors to promote GBM growth. *In vivo*, GR KD tumors grew strikingly less, by 5- to 6-fold, than WT tumors (Figure 3B, G) and these mice survived longer and lost less weight from start to end of the experiment (Figure 3C, H). As an independent assessment of tumor progression *in vivo*, we measured expression of a constitutive reporter *Ef1a*:luc in GBM xenografts. We found *Ef1a*-driven bioluminescence in LN229 and GL261 cells increased with cell number, without circadian modulation, *in vitro* (Figure S8A, C), and *in vivo* (Figure S8B, D). We next generated GR KD lines of LN229- and GL261-*Ef1a*:luc cells and tracked tumor size following implantation into the basal ganglia. Consistent with our previous findings, GR KD cells grew significantly less, by 2- to 5-fold, than WT xenografts (Figure 3D, I). These mice lost less weight from start to end of the experiment, indicating slower disease progression (Figure 3E, J). Finally, we measured Ki67 expression as an additional marker of cell proliferation and found significantly higher Ki67 expression and tumor area in WT tumor slices, compared to GR KD (Figure S9A-B). These results indicate that GR expression in GBM cells is required for endogenous glucocorticoids to promote tumor growth and disease progression.

**Figure 3:**
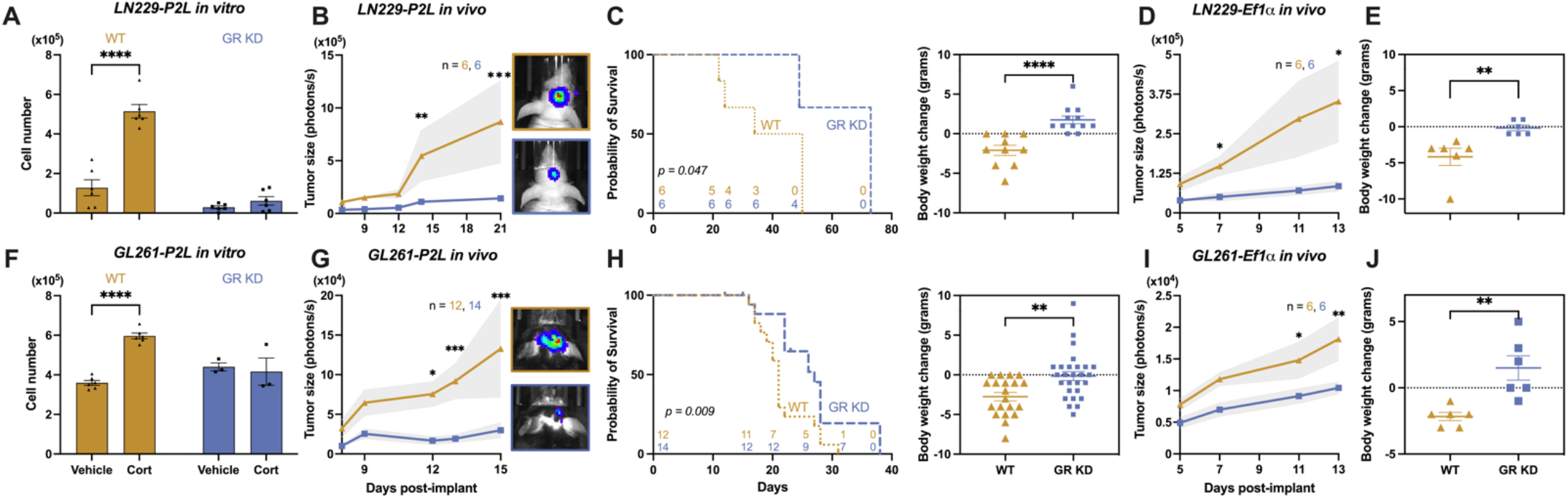
Daily glucocorticoid signaling promotes GBM growth and accelerates disease progression (A, F) Glucocorticoids promote cell growth *in vitro* in LN229 and GL261 WT, but not in GR KD cells (n=6 per condition, mean±SEM, n = 6 per group, two-way ANOVA with Bonferroni’s multiple comparisons test, ****p < 0.0001, ns p > 0.05). Cells treated with 100μM cortisol (LN229) or corticosterone (GL261) grew an average of 4- to 2-fold compared to vehicle-treated or GR KD cultures. See also Figure S7. (B, G) Tumor size is higher in mice bearing LN229 and GL261 WT tumors, compared to GR KD (mean±SEM, n reported in figure, two-way ANOVA with Šídák’s multiple comparisons test, *p < 0.05, **p < 0.01, ***p < 0.001). (C, H) Probability of survival is higher in mice bearing LN229 and GL261 GR KD tumors, compared to those implanted with WT tumors (*p<0.05). Mice bearing LN229 or GL261 GR KD tumors lost less weight from start to the end of the experiment, compared to mice implanted with WT tumors (mean±SEM, n reported in figure, t test, **p < 0.01, ****p < 0.0001). (D, I) Independent measurements of tumor bioluminescence using a constitutive *Ef1a*-luc reporter in LN229 and GL261 tumors shows higher tumor size in mice bearing WT tumors, compared to GR KD (mean±SEM, n reported in figure, two-way ANOVA with Šídák’s multiple comparisons test, *p < 0.05, **p < 0.01). See also Figure S8 and S9. (E, J) Mice bearing LN299 and GL261-*Ef1a* GR KD tumors lost less weight from start to the end of the experiment, compared to mice implanted with WT tumors (mean±SEM, n reported in figure, t test, **p < 0.01).

To further evaluate if daily rhythms in glucocorticoid secretion regulate GBM progression, we implanted GL261-P2L cells into the basal ganglia of mice with impaired circadian rhythms. We chose to study mice with the vasoactive intestinal peptide (VIP) gene knocked out because they lose circadian regulation of rest-wake activity and corticosterone secretion^39,47^. We measured fecal corticosterone every 4 hours for 24h in WT and VIP KO mice bearing GL261-P2L tumors and housed in constant darkness^29^. We found corticosterone secretion peaked during the subjective night (CT 12) in WT, but remained chronically low in VIP KO mice (Figure 4A). GBM tumors grew more in WT mice, by 11-fold, compared to VIP KO (Figure 4B). Furthermore, VIP KO mice bearing GL261-P2L tumors gained body weight from start to end of the experiment, whereas WT mice lost weight (Figure 4C), demonstrating that circadian signaling is required for aggressive GBM progression. Finally, we found significantly higher Ki67 expression and tumor area in tumors implanted into WT mice, compared to VIP KO (Figure 4D). Altogether, these results suggest that the daily surge in glucocorticoid secretion around waking promotes GBM growth and accelerates disease progression.

**Figure 4:**
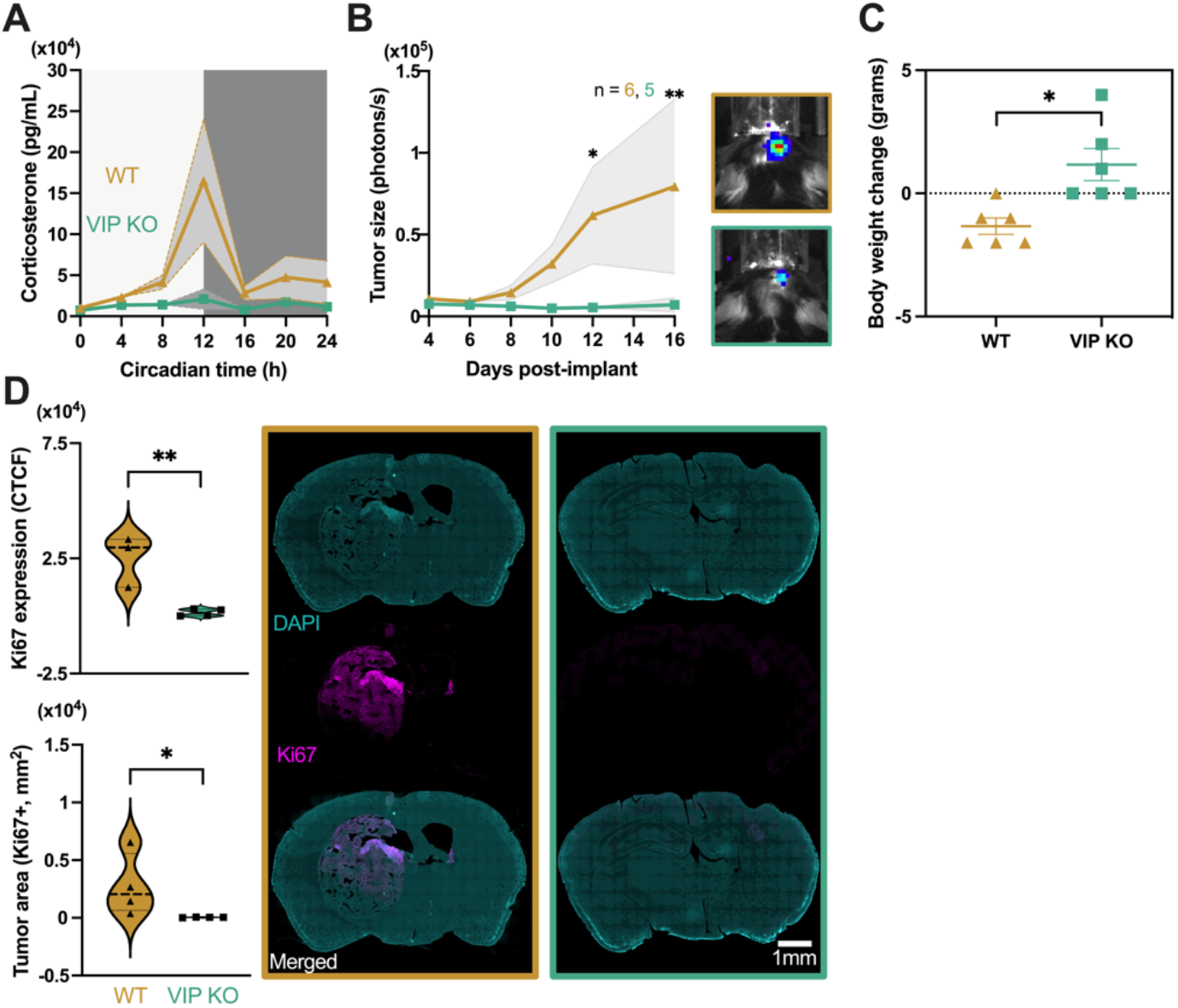
Disruption of circadian rhythms in the host slows GBM growth and disease progression (A) Fecal Corticosterone (CORT) concentration is rhythmic in WT, but not VIP KO mice (mean±SEM, WT trace scored circadian by JTK cycle p < 0.05). (B) Tumor size is higher in WT mice bearing GL261 tumors, compared to VIP KO mice (mean±SEM, n reported in figure, two-way ANOVA with Šídák’s multiple comparisons test, *p < 0.05, **p < 0.01). (C) VIP KO mice bearing GL261 tumors lost less weight from start to the end of the experiment, compared to WT mice (mean±SEM, n reported in figure, t test, *p < 0.05). (D) GBM proliferation and tumor area, as measured by Ki67 expression, is higher in WT mice bearing GL261 tumors, compared to VIP KO mice (mean±SEM, scale bar = 1mm, n = 4 per group, t test, *p < 0.05, **p < 0.01). DAPI (cyan) staining was used to label nuclei in whole brain sections. Composite images of DAPI and Ki67 (magenta) staining are shown to visualize tumor in the brain.

### Intrinsic daily rhythms of mouse and human GBM cells synchronize to the host’s central clock

To assess if GBM cells implanted into the basal ganglia act as circadian pacemakers that entrain to daily local cues, we recorded *Bmal1* expression from LN229 xenografts and running wheel activity of mice 2 weeks after reversing the 12 h:12 h light:dark cycle (LD to DL, lights on 7pm to 7am; Figure 5A). We found that, before the shift in the light cycle, tumor *Bmal1* expression peaked around ZT4 (8 h before daily locomotor activity onset) and *Per2* peaked around ZT18 (6 h after activity onset) in individual mice and group averages (Figure 5B, E). After 2 weeks in the reversed light schedule, tumor daily rhythms showed peak *Bmal1* around ZT0 and *Per2* around ZT13 (Figure 5C, F). We then evaluated these mice after 2 weeks in constant darkness (DD) and found that tumor rhythms shifted with the free-running period of locomotor activity of each mouse (*Bmal1* peaked at CT2 and *Per2* peaked at CT14; Figure 5D, G). These findings suggest that clock gene expression in GBM tumors entrains to the host’s central clock, shifting as mice entrain to changes in the light cycle.

**Figure 5:**
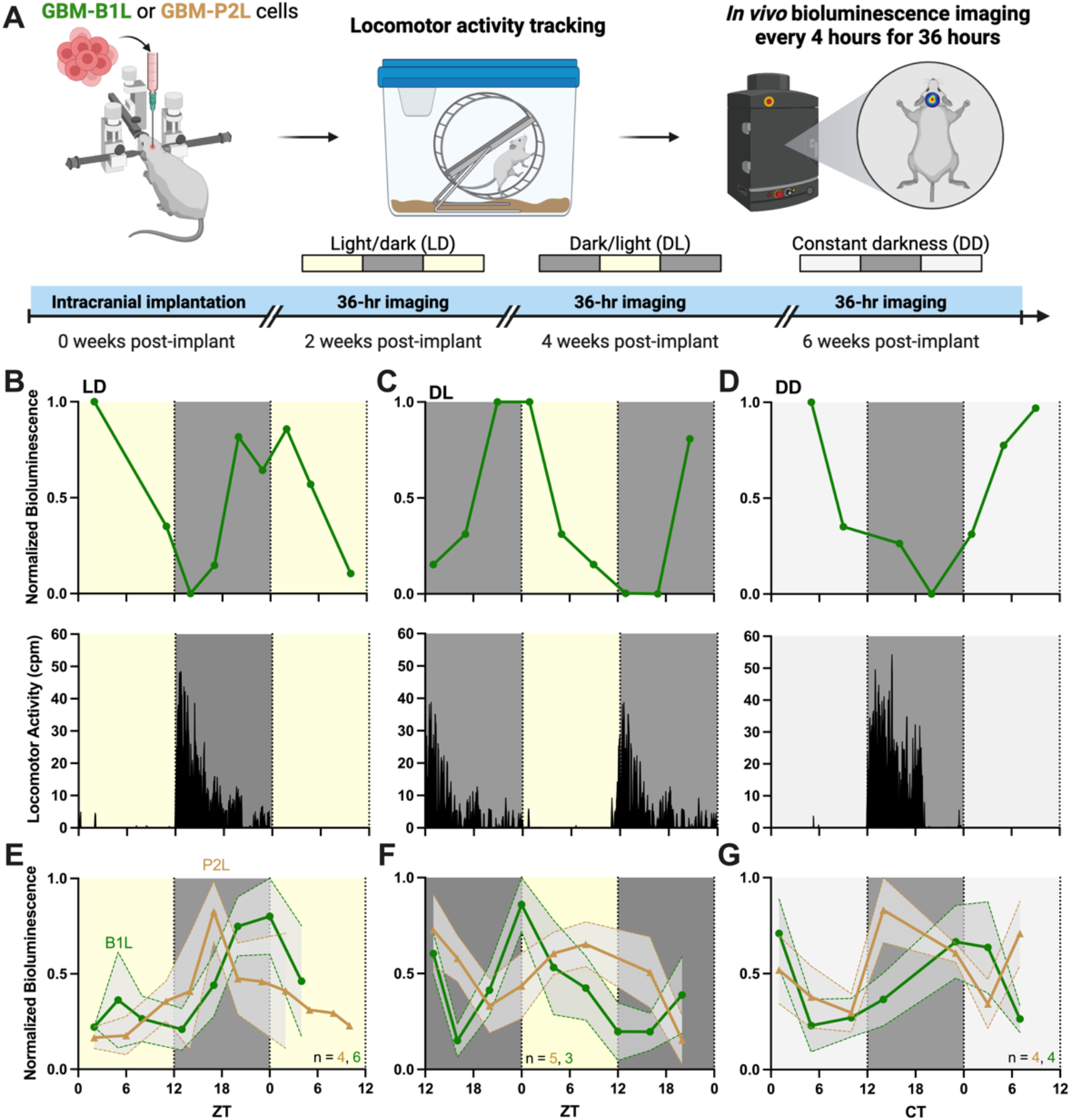
Peak timing of tumor *Bmal1* and *Per2* synchronizes to host rest-wake activity in cycles of light-dark, dark-light, and constant darkness (A) Schematic of light shifting paradigm after tumor implantation. (B) Representative 36-hour *in vivo* bioluminescence imaging (top) and locomotor activity profile (bottom) of a mouse implanted with NF1^-/-^ DNp53-B1L cells, in a standard 12L:12D light schedule, two weeks post-implant, shows peak *Bmal1* expression during the light phase (trace scored circadian by JTK cycle p < 0.05) and entrainment to the light cycle. (C) 36-hour *in vivo* imaging (top) and locomotor activity profile (bottom) of the same mouse after four weeks in a reversed 12L:12D light schedule, six weeks post-implant, shows *Bmal1* synchronized to the new dark-light cycle, peaking during the light phase (trace scored circadian by JTK cycle p < 0.05), and entrainment to the new light cycle. (D) 36-hour *in vivo* imaging (top) and locomotor activity profile (bottom) of the same mouse after four weeks in constant darkness, eight weeks post-implant, shows earlier *Bmal1* peak timing associated with the daily advancing of onsets observed in free running actigraphy (trace scored circadian by JTK cycle p < 0.05). Intrinsic locomotor started during the subjective night (CT12), showing the intrinsic circadian rhythm in locomotion. (E) Average traces of 36-hour *in vivo* imaging of NF1^-/-^DNp53 B1L and P2L GBM tumors two weeks post-implant, in a standard 12L:12D light schedule, show peak *Bmal1* during the light phase and *Per2* during the dark phase (mean±SEM, average traces scored circadian by JTK cycle p < 0.05). (F) Average traces of 36-hour *in vivo* imaging of NF1^-/-^ DNp53 B1L and P2L GBM tumors four weeks post-implant, after two weeks in a reversed 12L:12D light schedule, show synchronized tumor rhythms to the new light-dark schedule (mean±SEM, average traces scored circadian by JTK cycle p < 0.05). (G) Average traces of 36-hour *in vivo* imaging of NF1-/- DNp53 B1L and P2L GBM tumors six weeks post-implant, after two weeks in constant darkness, showing shifted peak timing (mean±SEM, average traces scored circadian by JTK cycle p < 0.05).

To further evaluate how daily rhythms in brain tumors entrain to the host, we implanted GL261-P2L GBM cells into the basal ganglia of WT and VIP KO mice, and recorded locomotor activity and GBM clock gene expression. We found that, in individual WT mice, *Per2* reliably peaked at night (CT 16), about 4 h after daily locomotor onset, but peaked at random times in individual VIP KO mice which lacked daily rhythms in locomotion (Figures 6A-B and S10A-H). Thus, group averaged *Per2* expression was circadian in WT (Figures 6 and S10), but not in VIP KO mice (Figures 6C-D and S10I-J). We used the time of daily peak *Per2* expression from the GBM xenograft of each mouse to quantify the high synchrony among tumors in WT, but not VIP KO mice (Figure S10K). Altogether, our findings indicate daily rhythms in the host are required to synchronize tumor daily *Per2* expression. We conclude that GBM tumors act as peripheral circadian oscillators that synchronize to the host’s central clock.

**Figure 6:**
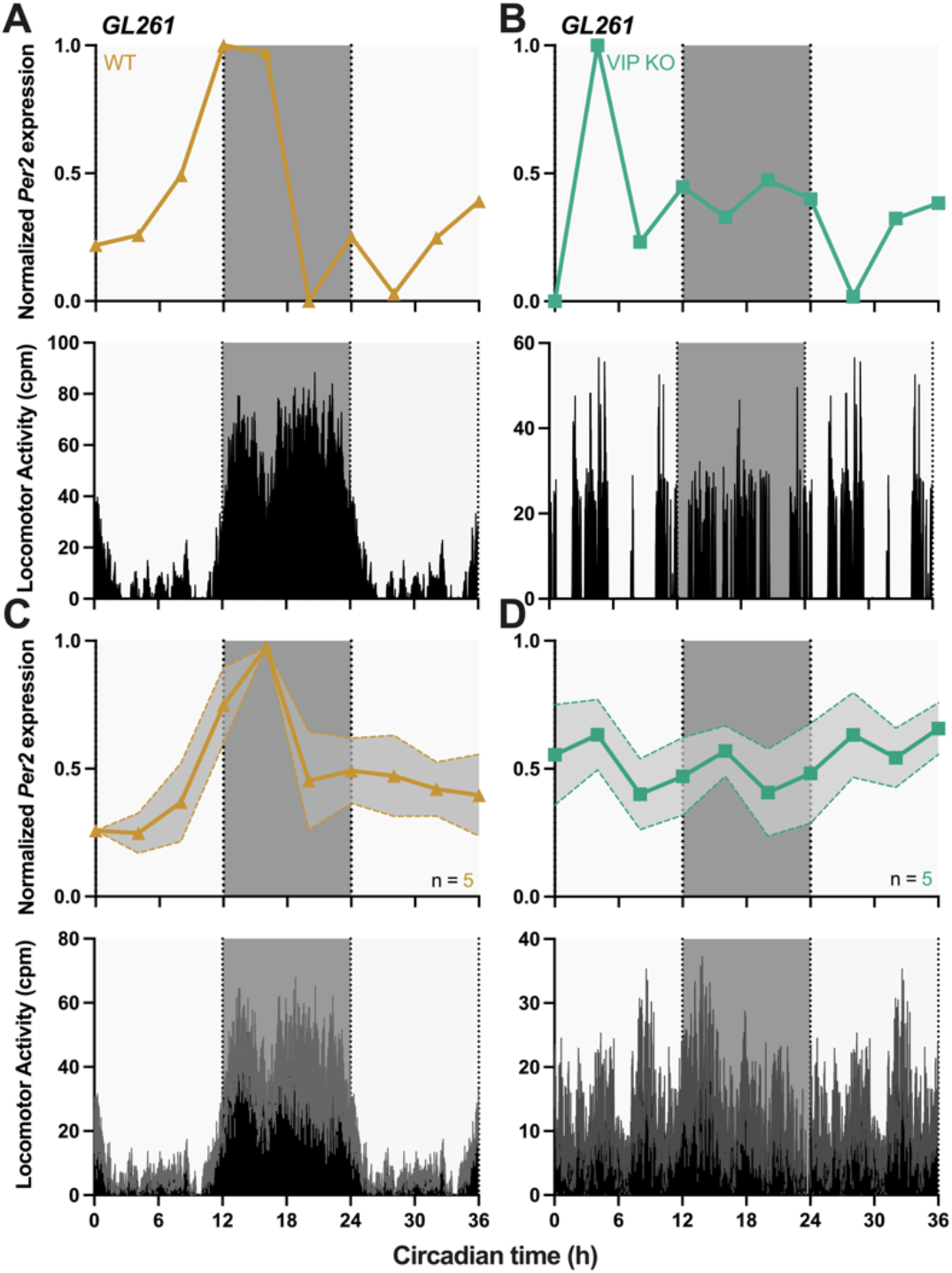
Disruption of daily rhythms in the host desynchronizes tumor *Per2* expression from the host rest-wake activity (A) Representative 36-hour *in vivo* bioluminescence imaging (top) and locomotor activity profile (bottom) of a WT mouse implanted with GL261-P2L cells, two weeks post-implant and in constant darkness, shows tumor *Per2* expression peaking during the subjective dark phase (CT16, trace scored circadian by JTK cycle p < 0.05) and rhythmic locomotor activity starting at CT12. (B) Representative 36-hour *in vivo* bioluminescence imaging (top) and locomotor activity profile (bottom) of a VIP KO mouse implanted with GL261-P2L cells, two weeks post-implant and in constant darkness, shows tumor *Per2* expression peaking during the subjective light phase (CT4, trace scored circadian by JTK cycle p < 0.05) and arrhythmic locomotor activity patterns. (C) Average trace of 36-hour *in vivo* imaging of WT mice (top) and average locomotor activity profiles (bottom) for all mice implanted with GL261-P2L tumors, two weeks post-implant and in constant darkness, shows reliable *Per2* peak expression and rhythmic daily activity (mean±SEM, average trace scored circadian by JTK cycle p < 0.05). (D) Average trace of 36-hour *in vivo* imaging of VIP KO mice (top) and average locomotor activity profiles (bottom) for all mice implanted with GL261-P2L tumors, two weeks post-implant and in constant darkness, shows desynchronized *Per2* peak timing and arrhythmic daily activity (mean±SEM, average trace not scored circadian by JTK cycle p > 0.05). See also Figure S10.

### GBM requires glucocorticoid receptors to synchronize daily rhythms to the host

Cell-autonomous circadian clocks have been found in brain regions outside the central pacemaker in the SCN, including the olfactory bulb, pineal gland, and pituitary^48–50^, as well as outside the brain, in the liver, kidney, lung, brown fat, heart, adrenal gland, among others^51–55^. These peripheral oscillators likely integrate signals from the central circadian clock in the SCN to regulate daily rhythms in cellular processes. Because glucocorticoids are known to be one of the most potent synchronizing cues for circadian clocks in peripheral tissues and we found that tumors fail to entrain normally in VIP KO mice with blunted sleep-wake and glucocorticoid rhythms, we hypothesized that daily glucocorticoids synchronize daily rhythms in GBM to the host. To test this hypothesis, we recorded *Per2* expression *in vitro* from LN229- and GL261-P2L cells, with or without GR KD, treated with 100nM DEX. We found that DEX rapidly induced *Per2* expression and shifted the daily rhythms in WT, but not GR KD, cells (Figure 7A-B, E-F). On the second day after treatment, DEX delayed *Per2* peak expression in LN229-P2L cells by 4 h, compared to vehicle (0.001% Ethanol), and by 5 h in GL261-P2L. Because DEX has a long half-life and thus can have lasting effects on daily gene expression, we next tested whether daily glucocorticoids can entrain GBM cultures. We recorded *Per2* expression from LN229 and GL261 cells *in vitro* and found that addition of glucocorticoids (100μM cortisol or corticosterone) at CT0 for 3 consecutive days synchronized the time of daily peak *Per2* expression to a 24 h rhythm compared to vehicle (0.001% ethanol; Figure S11A-D). Cultures resumed their free-running circadian periods after termination of daily glucocorticoid administration. Together, these findings suggest that daily glucocorticoid treatment entrains intrinsic *Per2* expression in GBM *in vitro*.

**Figure 7:**
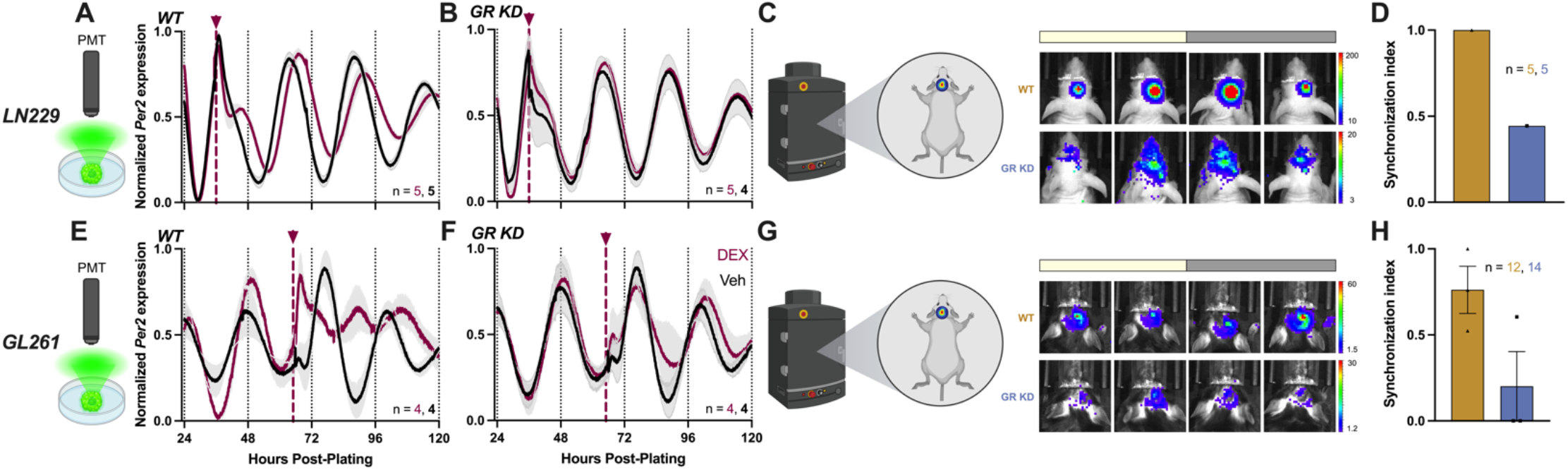
Timing of daily *Per2* expression in GBM xenografts depends on glucocorticoid receptor signaling (A, E) Acute addition of 100nM Dexamethasone induces a phase shift in circadian *Per2* expression, compared to vehicle, in WT LN229 and GL261 GBM cells (mean±SEM, all recordings had cosine fits with correlation coefficients, CC > 0.9). See also Figure S11. (B, F) The glucocorticoid receptor is required for Dexamethasone-induced phase shifting of *Per2* expression in LN229 and GL261 GBM cells (mean±SEM, all recordings had cosine fits with correlation coefficients, CC > 0.9). (C, G) Representative images of *in vivo* tumor imaging during the light (ZT0-12) and dark (ZT12-24) phases of LN229-P2L and GL261-P2L, WT or GR KD cells. (D, H) *Per2* expression is highly synchronized in LN229 and GL261 WT tumors, but desynchronizes in tumors lacking GR (mean±SEM). See also Figure S12.

To test if GR signaling coordinates GBM circadian rhythms *in vivo*, we next recorded clock gene expression from GR-deficient xenografts in tumor-bearing mice. We implanted WT or GR KD GBM cells into the basal ganglia of mice (LN229-P2L into immunocompetent nude or GL261-P2L into C57 WT). After recovery and detection of bioluminescence over background, we monitored *Per2*-driven bioluminescence every 4 h for 36 h (Figure 7C, G). We found tumor *Per2* expression reliably peaked in the middle of the night (ZT 12-20) in LN229 and GL261 WT cells, but peaked at varying times of day in GR-deficient GBM cells, as evidenced by a lower synchronization index among mice and misalignment with the daily locomotor activity onset (Figure 7C-D, G-H and S12A-L). We conclude that timing of daily clock gene expression in GBM depends on glucocorticoid receptor signaling *in vivo*.

## Discussion

### Chronotherapy using Dexamethasone for GBM

The effects of DEX on GBM progression have remained controversial due to studies demonstrating anti- and pro-proliferative effects, depending on cell type, drug concentration and experimental conditions. Strikingly, many of these studies used the same GBM models and drug concentration, but obtained opposite effects. We identified circadian time as an important variable explaining the differential tumorigenic effects of DEX. We found a 3-fold increase in GBM cell growth when delivering DEX around the daily peak of tumor *Bmal1* expression, and a 3-fold growth suppression around the trough. Further, we found that DEX-induced tumor growth depends on the core clock gene *Bmal1*. In mice bearing GBM xenografts, DEX delivered during the morning significantly increased GL261 growth by 2-fold and LN229 growth by 5-fold, compared to those treated in the evening or with vehicle. Tumor-bearing mice receiving DEX in the morning lost more weight from start to end of the experiment, compared to evening or vehicle groups, indicating accelerated disease progression. These results suggest circadian regulation of the tumor’s response to the synthetic glucocorticoid and represents a striking example of how chronotherapy (when a drug is delivered relative to circadian time) differentially affects cancer outcomes.

DEX mimics the effects of endogenous glucocorticoids, a hormone secreted in a circadian fashion, by binding to glucocorticoid receptors and activating this signaling pathway^56,57^. Thus, DEX action on glucocorticoid receptor signaling may be intrinsically regulated by the circadian clock, and its effects may vary with time of day of drug administration. Our results indicate that DEX-induced proliferation depends on *Bmal1* which may relate to recent findings on glucocorticoid and circadian regulation of the cell cycle^58–60^. For example, a recent study found that DEX promotes GBM growth by down-regulating genes controlling G2/M and mitotic-spindle checkpoints, and up-regulating anti-apoptotic regulators BCL2L1 and MCL1^58^. DEX binding to GR may also be circadian leading to rhythms in activation of glucocorticoid response elements (GREs) and transactivation or transrepression of transcription. Future studies should test how the circadian clock modulates glucocorticoid action in GBM.

Our findings have important implications for the use of DEX in GBM patients. Recent findings have demonstrated the benefits of timing therapies (i.e., chronotherapy) to maximize chemotherapy efficacy and tumor death in GBM^6–8,32^. While it hasn’t been studied for GBM, chronotherapy using corticosteroids has yielded positive effects for diseases like multiple sclerosis and rheumatoid arthritis^61,62^. Implementing a chronotherapeutic approach for DEX use in the clinic may maximize its anti-inflammatory benefits while minimizing its effects on GBM growth. Further, it will be important to consider whether timing other corticosteroids with shorter half-life yields similar effects as DEX, reducing GBM-induced edema while protecting from tumor growth. If validated in patients, incorporating chronotherapy into the standard of care for DEX use in GBM patients requires no additional approvals or clinical trials. Future studies should focus on evaluating whether the time at which DEX administration protects from GBM growth also yields a reduction in cerebral edema. Moreover, because DEX has a long half-life of 36-72 hours, it will be important to consider whether timing its administration in the clinic yields anti- or pro-proliferative effects, and whether other corticosteroids may represent better options for GBM patients.

### The clock is ticking in GBM: Intrinsic daily rhythms synchronize to the host through glucocorticoids

Our findings support a model where GBM tumors act as peripheral circadian oscillators that can integrate into circadian circuits of the brain. This has implications for diagnosis and treatment. In the two human and two mouse GBM models tested, regardless of sex or immune status of the host, *Bmal1* expression peaked during the day and *Per2* peaked at night, suggesting a conserved mechanism that synchronizes circadian rhythms across a variety of GBM cell types and genotypes. Recent findings in humans, cellular, and animal models of GBM have shown increased sensitivity to chemotherapy with TMZ when delivered in the morning^6,8^. This can now be explained based on the diversity of GBM models in this study that similarly entrain to the local light cycle in mice. This leads to the hypothesis that measuring daily rhythms in the host (e.g., sleep-wake or cortisol) could be used to guide optimized time of treatment for GBM.

How do brain cancers synchronize their daily rhythms to the host? We find, for example, that circadian gene expression in GBM xenografts adjusts to changes in the light cycle, synchronizes to daily rest-activity patterns, and depends on the neuropeptide, VIP, and daily hormones, such as glucocorticoids. One possibility is that SCN-driven signals dictate the internal phase of clock gene expression by acting as timing cues to peripheral oscillators. Multiple signaling pathways, such as calcium, cAMP, protein kinase A and C, vasoactive intestinal peptide, glucocorticoids, and others, have been implicated in the regulation of phases of peripheral oscillators37,38,42,44,47,63–70.

Among mammalian peripheral tissues, manipulation of glucocorticoid signaling resets the phase of clock gene expression in liver, heart, and kidney^42^. For example, the glucocorticoid receptor agonist Dexamethasone induces *Per1* gene expression in liver, kidney, and rat primary hepatocytes, and down-regulates *Reverbα* in liver tissues^42^. Here we found that Dexamethasone treatment shifts the phase of *Per2* gene expression in human and murine GBM cells *in vitro*, dependent on intact glucocorticoid receptor signaling. We present evidence that reveals that daily glucocorticoids synchronize daily rhythms in GBM tumors to the host, maintaining reliable clock gene expression patterns throughout the day. In the absence of glucocorticoid receptors or by disruption of glucocorticoid rhythms (i.e., VIP KO), GBM tumors sustain their daily rhythms, but *Per2* gene expression peaks at random times of day, suggesting that these peripheral signal acts as a time giver for the tumor. Our findings support a model whereby the master clock in the SCN regulates daily rhythms in glucocorticoid secretion that then synchronize daily clock gene expression in GBM tumors. Future studies should explore the mechanisms underlying how glucocorticoids and other daily signals synchronize daily rhythms in GBM to the host.

### Daily rhythms in glucocorticoid secretion regulate GBM progression

Previous research has centered on whether synthetic glucocorticoids like DEX regulate GBM progression, yet no studies have considered the role of daily rhythms in endogenous glucocorticoid signaling. Our findings identify daily rhythms in glucocorticoids as a regulator of GBM progression. We found smaller tumors, longer host survival, and less body weight loss in mice bearing GBM tumors that lack the glucocorticoid receptor (i.e., GR KD). Tumor size decreased by 5- to 6-fold when lacking GR, compared to WT tumors. Further, tumors implanted into mice that lack a daily surge in glucocorticoids (i.e., VIP KO) grew significantly slower than tumors in WT mice by 11-fold. Altogether, we propose that the daily surge in glucocorticoid secretion promotes GBM growth and accelerates disease progression.

The finding that daily glucocorticoid signaling promotes GBM growth opens the doors to a deeper mechanistic understanding of its role in glioma tumorigenesis. Glucocorticoids maintain various metabolic and homeostatic functions by binding to the glucocorticoid receptor. In the absence of glucocorticoids, GR is localized to the cytoplasm, bound to chaperone proteins such as HSP90. Upon ligand binding, GR undergoes a confirmational change that triggers its translocation to the nucleus, where it can exert its actions primarily through genomic transactivation and transrepression, or by non-genomic signaling mechanisms^57^. The mechanisms by which this signaling program is regulated by the circadian clock remain poorly understood. Beyond regulating normal tissue function, the role of GR in cancer has been understudied beyond murine lymphoma, human leukemia cells, mouse primary thymocytes, human primary chronic lymphoblastic leukemia, and acute lymphoblastic leukemia, in which GR acts as a proapoptotic cue^71–75^. In other cell types, however, GR has been shown to coordinate cell division and mitosis, with loss of GR leading to aberrant chromosome segregation, accumulation of chromosome complement defects, and a robust mitotic phenotype^76^. While the role of GR in GBM has received very little attention, our findings suggest that this signaling pathway regulates tumor progression by inducing a proliferative phenotype, potentially depending on circadian clock regulation of GR signaling. These findings highlight an interaction between circadian circuits and GBM, and identifies a targetable mechanism regulating GBM progression.

Beyond maintaining synchronized daily rhythms between the tumor and the host, the circadian clock has been implicated in GBM tumorigenesis. For example, the CLOCK-BMAL1 complex promotes tumor angiogenesis and growth in GBM^77,78^. Disrupting the circadian clock by downregulating BMAL1 and CLOCK induces cell-cycle arrest and apoptosis in GBM stem cells^31^. In humans, analysis of TCGA data revealed that high expression of BMAL1 correlated with shorter patient survival times^31^. Altogether, these data led us to ask whether synchronized circadian rhythms drive GBM progression. We found that desynchronizing daily rhythms in GBM tumors from the host’s clock by disrupting glucocorticoid receptor signaling significantly reduced tumor growth and slowed disease progression *in vivo*. Strikingly, tumors implanted into arrhythmic VIP KO mice, which also desynchronize from the host’s central clock, fail to grow compared to tumors implanted into WT mice. Together, our findings suggest that synchronized circadian rhythms through glucocorticoid receptor signaling drive GBM progression.

Because of the localization of GBM tumors and their sensitivity to respond to peripheral signals, we do not exclude the possibility of other neural or non-neuronal signals acting as timing cues. Further, it is possible that timing cues also regulate glioma progression dependent on time of day. A wealth of recent data has presented evidence for a reciprocal crosstalk between gliomas and neurons in the tumor microenvironment that drives tumor progression^79–85^. Neuronal activity has been found to promote glioma growth^82,83,85^, while gliomas have been found to remodel adjacent neuronal synapses to induce a state of brain hyperactivity^79,81,83,84^. This raises the possibility that neuron-GBM communication synchronizes circadian rhythms in these tumors and promotes growth at specific times of day. Because neuronal activity and secretion of mitogens (i.e., Neuroligin-3, Semaphorin-4F, EphrinA6, and EphrinA7) varies with time of day^80,86–90^, it is possible that activity-driven glioma growth depends on circadian time. Outside of the brain, temperature, and Insulin/Insulin-like growth factor (IGF) in combination with glucose can act as timing cues for many tissues^40,41,44^. Whether these signals promote daily GBM progression has not been studied. Future studies could test the role of brain temperature rhythms, and daily food intake and metabolism in tumor entrainment and growth. Moreover, future studies could focus on whether GBM is more sensitive to promitotic cues at specific times of day, if SCN and clock-derived signals regulate glioma progression, and whether neuron-GBM reciprocal crosstalk depends on circadian time. Altogether, this line of research will provide insights into how gliomas integrate into brain circuits, hijack its normal physiology, and use daily cues to grow. These findings could be leveraged to identify novel therapeutic targets that can slow GBM progression, while opening avenues for treatment optimization by delivering therapies in accordance with a tumor’s circadian rhythm.

### Beyond GBM: targeting the clock in cancer biology

Chronotherapy, also referred to as circadian medicine or chronomedicine, seeks to treat patients at the optimal time of day to maximize health benefits and minimize side effects. This approach has been shown to be effective in acute lymphoblastic leukemia^91,92^, colorectal^93^, ovarian, other gynecological cancers^94^, and most recently GBM^6,8^, but is neither the standard of care nor carefully evaluated in many cancers. We recently demonstrated that GBM cells and xenografts are more sensitivity to chemotherapy with TMZ when delivered at the daily peak of *Bmal1* clock gene expression *in vitro* and in the morning *in vivo*^8^. Daily sensitivity to TMZ is regulated by circadian expression of the DNA repair enzyme MGMT, and inhibiting its expression abolishes the daily rhythm in sensitivity to the drug^8^. In humans, timing TMZ to the morning extended patient survival by 6 months, compared to administering chemotherapy in the evening^6^. Here we found that, unlike some cancer types, GBM sustains intrinsic circadian rhythms that synchronize to the host and use daily clock-controlled cues to grow. Further, these daily rhythms can be leveraged to identify the best times of day to treat GBM tumors with drugs like TMZ and DEX.

Chronotherapy also considers daily signals, such as light in the morning and melatonin before sleep, as cues that synchronize a patient’s daily rhythm. Previous findings show that tumor progression can be slowed by enhancing or inducing circadian rhythmicity in the host or in cancer tissues, such as breast^95^, melanoma^96^, colon carcinoma^96^, osteosarcoma^97^, and pancreatic adenocarcinoma^98^. To critically evaluate the potential for chronotherapy in different cancers, we must consider how daily rhythms arise and synchronize in specific tissues. Given that about 80% of approved drugs in the United States hit targets known to rise and fall according to time of day and intrinsic circadian rhythms^99,100^, it is possible that many therapeutics used to treat different cancers work best at specific times of day. It will be important to understand how circadian rhythms regulate tumor biology in a cell- and tissue-specific context. Altogether, this highly tractable and translatable approach can ultimately personalize patient care by determining when therapies should be given to cancer patients, depending on their individual circadian rhythms.

## Acknowledgments

The authors thank the members of the Herzog lab for valuable discussions and comments on the manuscript. This work was supported by National Institutes of Health Grants NINDS R21NS120003, NCI R01NS134885, and the Washington University Siteman Cancer Center. Author MFGA was supported by the Washington University Neuroscience Program T32-Training Grant NIH (T32NS121881-01) and the Initiative for Maximizing Student Development (IMSD) Program Training Grant NIH (R25GM103757-10). Author ARD was supported by the National Institutes of Health National Cancer Institute (F31CA250161).

## Author contributions

All authors contributed to the study conception and design. Cell experiments and analysis: MFGA, ARD, TS, and FD. Animal experiments and analysis: MFGA and ARD. ELISA and tissue processing: MFGA and NA. The first draft of the manuscript was written by MFGA and EDH. All authors read and approved the final manuscript.

## STAR Methods

### RESOURCE AVAILABILITY

#### Lead contact

Further information and requests for resources and reagents should be directed to and will be fulfilled by the lead contact, Dr. Erik Herzog.

#### Materials availability

This study did not generate new unique reagents.

#### Data and code availability

- All data reported in this paper will be shared by the lead contact upon request.
- This paper does not report original code.
- Any additional information required to reanalyze the data reported in this paper is available from the lead contact upon request.

## EXPERIMENTAL MODEL AND STUDY PARTICIPANT DETAILS

### Glioblastoma cell culture

#### GL261

Glioma 261 (GL261, obtained from the Division of Cancer Treatment and Diagnosis Tumor Repository of the National Cancer Institute), a male murine model of GBM, were cultured in monolayer in coated T-75 flasks (Nunc Treated EasYFlasks, Fisher) using RPMI-1640 media (Sigma-Aldrich), supplemented with 10% FBS (Fisher), 1% L-Glutamine (Thermo Fisher), and 1% Pen/Strep (Thermo Fisher). Cells were grown in a 37°C incubator with a 5% CO2 environment. Passage number in all experiments ranged from four to eight.

#### NF1^-/-^DNp53

Nf1^-/-^DNp53 male astrocytes (generous gift of Dr. Josh Rubin), a murine model of GBM (Warrington et al., 2007), were cultured in monolayer in coated T-75 flasks (Nunc Treated EasYFlasks, Fisher) using 10mL DMEM/F12 media (Gibco), supplemented with 10% FBS (Fisher) and 1% Pen/Strep (Thermo Fisher). Cells were grown in a 37°C incubator with a 5% CO2 environment. Passage number in all experiments ranged from four to eight.

#### B165

Human B165 (MGMT methylated, male), generous gift of Dr. Albert Kim, were cultured as spheres in 100mm uncoated petri dishes (Fisher) using 3mL DMEM/F12 Glutamax media (Gibco), supplemented with 1% Pen/Strep (Thermo Fisher), 2% B-27 (Miltenyi Biotec), 25ug/10mL Heparin (Sigma-Aldrich), 20ug/100ml EGF (Pepro Tech), and 2ug/100ml bFGF (Pepro Tech). Cells were grown in a 37°C incubator with a 5% CO2 environment. Passage number in all experiments ranged from six to ten.

#### LN229

LN229 (American Type Culture Collection), a female human cell line, were cultured in monolayer in coated T-75 flasks (Nunc Treated EasYFlasks, Fisher) using 10mL DMEM media (Gibco), supplemented with 5% FBS (Fisher), and 1% Pen/Strep (Thermo Fisher). Cells were grown in a 37°C incubator with a 5% CO2 environment. Passage number at implant ranged from four to eight.

### Animals

#### C57BL/6NJ

10-week-old immunocompetent male and female C57BL/6NJ (The Jackson Laboratory, Strain #000664) mice were used in all experiments for orthotopic xenografting of murine GBM cells. Mice were singly housed in standard 12 h light/12 h dark conditions in individual wheel-cages to record locomotor activity in one-minute bins. After surgery, mice were monitored and treated with analgesic for three days. For some experiments, light schedules were shifted to a reversed 12h:12h dark/light, or to constant darkness (DD). At the end of all experiments, or when subjects lost 10% body weight from start to end of the experiment, mice were sacrificed in accordance with IACUC protocols and brain tissue was collected for subsequent analysis.

#### CrTac:Ncr-Foxn1nu10

10-week-old immunocompromised male and female CrTac:Ncr-Foxn1nu (Taconic, Strain #NCRNU) mice were used in all experiments for orthotopic xenografting of human and primary GBM cells. Mice were singly housed in standard 12 h light/12 h dark conditions in individual wheel-cages to record locomotor activity in one-minute bins. After surgery, mice were monitored and treated with analgesic for three days. For some experiments, light schedules were shifted to a reversed 12h:12h dark/light, or to constant darkness (DD). At the end of all experiments, or when subjects lost 10% body weight from start to end of the experiment, mice were sacrificed in accordance with IACUC protocols and brain tissue was collected for subsequent analysis.

#### VIP KO

10-week-old immunocompetent male and female VIP KO (obtained from: Dr. James Waschek, UCLA) mice were used in experiments for orthotopic xenografting of human and primary GBM cells. Mice were singly housed in constant darkness conditions in individual wheel-cages to record locomotor activity in one-minute bins. After surgery, mice were monitored and treated with analgesic for three days. At the end of all experiments, or when subjects lost 10% body weight from start to end of the experiment, mice were sacrificed in accordance with IACUC protocols and brain tissue was collected for subsequent analysis.

## METHOD DETAILS

### Experimental cell culture

GBM cells were grown to confluence (passage no. < 10) in supplemented media conditions mentioned in the “Experimental model and study participant details” section, grown in a 37°C incubator with a 5% CO2 environment. Confluent cultures were kept for up to 4 weeks with media refreshed every 3-5 days. Once cultures reached 80% confluence, cells were split using TrypLE™ Express Enzyme (1X) (Gibco), centrifuged for 4 minutes at 0.4 RCF, diluted, and replated at a lower confluence in new cell culture flasks. Before the start of each experiment, cells were synchronized by changing culture media to a serum-free, supplemented with B27 (Thermo Fisher) “Air Media” (Bicarbonate-free, DMEM, 5mg/mL glucose, 0.35 mg/mL sodium bicarbonate, 0.01 M HEPES, 2 μg/mL pen/strep, 1% Glutamax, 1 mM luciferin, pH 7.4, 350 mOsm; adapted from Hastings et al., 2005), and dishes were sealed with sterile vacuum grease.

### Circadian luciferase lentivirus production

Lentiviral reporters expressing firefly luciferase driven by the mouse *Bmal1* (*Bmal1*-luc) (previously described in Liu et al., 2008; Zhang et al., 2009) or *Period2* (*Per2*-luc) (previously described in Ramanathan et al., 2012) promoters were generated in the laboratory using plasmids (gift from Dr. Andrew Liu, University of Memphis) and packaging plasmids psPAX2 and pMD2.G (gift from Didier Trono (Addgene plasmid # 12260 and 12259; RRID:Addgene_12260 and 12259)). Hek293T/17 cells (ATCC) were plated at 3 million cells per 100 mm dish in complete DMEM media with 10% FBS (Thermo Fisher) and 5% Pen/Strep (Thermo Fisher) in a 5% CO_2_ incubator for 24-30 hours. Before transfection, media was changed to DMEM with 5% FBS (Fisher) without any antibiotics. For transfection, cells were incubated with Fugene 6 (Promega) and plasmids for 12 hours, then changed to complete media. Viral particles in media were collected after 48 and 72 hours, centrifuged (RT, 3 min at 0.4 rcf) and filtered through a PES 0.45 um filter (Fisher). The presence of viral particles was confirmed with Lenti-X GoStix Plus test (Takara).

### Lentiviral transductions

#### Luciferase reporters for Bmal1, Per2, & Ef1a

GBM cells were transduced with lentiviral reporters expressing firefly luciferase driven by the mouse *Bmal1* (*Bmal1*-luc), *Period2* (*Per2*-luc), or *Ef1α* (*Ef1α-*luc, obtained from GenTarget Inc.). Cells were grown in T-25cm^2^ flasks for 24h and incubated for 10 minutes at 37°C in 3mL complete DMEM media with 10% FBS (Thermo Fisher), 5% Pen/Strep (Thermo Fisher), and 15μg polybrene (Millipore #TR-1003-G). Following incubation, 500uL of virus stock solution was added to each culture. Media was changed after 24 h at 37°C. Infected cells were selected using blasticidin (1.25 μg/mL, Thermo Fisher). Luciferase expression was confirmed by recording bioluminescence *in vitro*.

#### Glucocorticoid receptor knockdown

To knockdown *NR3C1* (GR) in LN229 and GL261 cells, two predesigned shRNA plasmids packaged into a lentivirus (vector pLKO.1) were obtained from Sigma Aldrich (Target sequences are listed in the Key resources table). Cells were grown in T-25cm^2^ flasks for 24h and incubated for 10 minutes at 37°C in 3mL complete DMEM media with 10% FBS (Thermo Fisher), 5% Pen/Strep (Thermo Fisher), and 15μg polybrene (Millipore #TR-1003-G). Following incubation, 10μL of virus stock solution was added to each culture. Media was changed after 24 h and cells were kept at 37°C, 5% CO2. Infected cells were selected using puromycin (1.5 μg/mL, Thermo Fisher) for 10 days. Knockdown efficiency was quantified by qPCR and immunocytochemistry. Two lentiviral constructs were independently tested.

#### Bmal1 knockdown

To knockdown *Bmal1* in LN229 cells, two predesigned shRNA plasmids packaged into a lentivirus (vector pLKO.1) were obtained from Sigma Aldrich (Target sequences are listed in the Key resources table). Cells were grown in T-25cm^2^ flasks for 24h and incubated for 10 minutes at 37°C in 3mL complete DMEM media with 10% FBS (Thermo Fisher), 5% Pen/Strep (Thermo Fisher), and 15μg polybrene (Millipore #TR-1003-G). Following incubation, 5-10μL of virus stock solution was added to each culture. Media was changed after 24 h and cells were kept at 37°C, 5% CO2. Infected cells were selected using puromycin (1.5 μg/mL, Thermo Fisher) for 10 days. Knockdown efficiency was quantified by qPCR and by recording *Per2*-luc bioluminescence *in vitro*. Two lentiviral constructs were independently tested.

### Bioluminescence recordings in vitro

Cells were plated in 35 mm (BD Falcon, Fisher) petri dishes, synchronized by a media change, supplemented with 1mL of serum-free, B27 containing “Air DMEM media” containing 0.1mM D-luciferin (Goldbio), sealed with vacuum grease, and placed in a light-tight 36°C incubator containing photo-multiplier tubes (PMTs) (Hamamatsu Photonics). Each dish was placed under one PMT and bioluminescence was recorded as photons per 180 seconds.

### Quantitative real-time PCR (qRT-PCR)

RNA was extracted from 500,000 cultured GBM cells using TRIzol reagent (Thermo Fisher). RNA was purified using the Direct-zol RNA MiniPrep Plus kit (Zymo) and cDNA was generated by RT-PCR using SuperScript® III First-Strand Synthesis System (Thermo Fisher). Gene expression changes were further probed using iTaqTM Universal SYBR® Green Supermix (Bio-Rad). The primer sequences for qRT-PCR are listed in the Key resources table. PCR amplification was carried out at 40 cycles with 100ng of template DNA in triplicates. Protocol is as follows: Cycle 1: 95 °C for 3 min; Cycle 2: 95 °C for 30 s; Cycle 3: 60 °C for 30 s; repeat step 2–3 for 39 more times; Cycle 4: 72 °C for 1 min. Negative controls included no reverse transcriptase reactions and no template DNA samples. All procedures were done in triplicate in two biological replicates.

### Immunocytochemistry

Cells were fixed using 4% PFA for 10 minutes, permeabilized for 30 min with 3% Triton-X (Millipore Sigma) in 1x PBS, and blocked for 1 hours with solution containing 10% BSA (Sigma) and 0.3% Triton-X. Primary antibodies were diluted in 2% blocking solution and incubated overnight at 4°C. Samples were then rinsed three times with 1x PBS and incubated in secondary antibody solution in 2% blocking solution for 1 hour at RT. Samples were rinsed 3 times in PBS, stained with ProLong Gold mounting medium with DAPI (Life Technologies, Carlsbad, CA), and stored in darkness at 4°C until imaging. Microscopy analysis was performed using ImageJ software. All antibodies and reagents used are listed in the Key Resources Table.

#### *In vitro* glucocorticoid entrainment assays

To assess whether glucocorticoids shift *Per2* expression *in vitro*, GBM-P2L cells were plated as previously described for bioluminescence recordings *in vitro*. After at least 24h of recording, cells were acutely treated with 100nM Dexamethasone or vehicle (0.001% Ethanol), and bioluminescence was recorded for at least 2 days post-treatment. Bioluminescence data was detrended with a 24-hour moving average and analyzed in ChronoStar 1.0.

To assess if glucocorticoids entrain *Per2* expression, GBM-P2L cells were plated as previously described for bioluminescence recordings *in vitro*. After at least 24 hours of recording, cells were treated with 100μM Corticosterone or Cortisol, for murine or human cells, respectively, or vehicle (0.001% Ethanol). Glucocorticoids were given chronically for 3 days at the same time of day. Bioluminescence was continuously recorded for at least 2 days post-treatment. Bioluminescence data was detrended with a 24-hour moving average and analyzed in ChronoStar 1.0.

#### *In vitro* cell growth assays and pharmacology

We calculated total cell number by labeling cells with DAPI and measuring fluorescence at 405nm. Cells were seeded in 6-well plates at the same cell density (100,000 cells/well) and kept on an incubator for 48h to allow for cell growth. To assess intrinsic growth of GBM cells, individual cell culture plates were fixed with cold methanol and stained with 2μg/mL DAPI at either 24, 48, or 72h post-plating. To assess whether glucocorticoids promote GBM growth, cells were treated 48h post-plating with 100nM Dexamethasone (for human and murine cells), 100μM Cortisol (only for human cells), 100μM Corticosterone (only for murine cells), or vehicle (0.001% Ethanol for DEX cultures; water for Cortisol or Corticosterone). To assess whether glucocorticoids promote GBM growth at specific times of day, cells were treated at two different time points. In all experiments, cells were fixed 72h post-treatment with cold methanol and stained with 4′,6-diamidino-2-phenylindole (DAPI, 2 μg/mL). DAPI fluorescence was quantified with the Infinite 200 PRO plate reader (V_3.37_07/12_Infinite, Tecan Lifesciences).

### Orthotopic xenografting

200,000 GBM cells were stereotactically implanted into the right caudate putamen (coordinates: Bregma, 2mm right laterally, 3mm ventral) of 10-week-old male and female NCr-Foxn1nu (Taconic Biosciences, Strain #NCRNU) for human models, C57BL/6NJ (The Jackson Laboratory, Strain #005304) for murine models, or VIP KO mice for murine models. After surgery, mice were housed in individual cages, monitored, and treated with analgesic for three days post-implant. Mice were allowed to recover and cells to engraft for 7 days before performing *in vivo* bioluminescence imaging to measure tumor size or daily rhythms in clock gene expression.

### In vivo bioluminescence imaging

Mice were housed in standard 12 h light/12 h dark, reversed 12h dark/ 12 h light, or constant darkness (DD) conditions in wheel-cages to record locomotor activity in one-minute bins. Following orthotopic xenografting, tumor size was consistently measured at 1:00 p.m. by anesthetizing mice with 2% isoflurane, subcutaneously injecting them with 15 mg/mL of D-luciferin, allowing 10 min for it to access the brain, and imaging bioluminescence with 5 min exposure time. To measure daily clock gene expression in GBM xenografts, imaging was performed as described every 4h for 36h. All imaging was performed using an *In Vivo* Imaging System Lumina III (IVIS, Perkin Elmer). Bioluminescence images were analyzed using Living Image Software (Perkin Elmer).

For long-term recordings of daily clock gene expression, mice were individually placed in constant darkness in a light tight box containing two photomultiplier tubes (PMT, Hamamatsu H8259-01, Lumicycle *In Vivo*, Actimetrics). Mice received luciferin in their water (10 mM, Gold Biotechnology) ad libitum over the 2 days of recording. This method allowed for recording of *Per2* or *Bmal1* luciferase expression in freely moving mice. Animals were checked daily to ensure that they had adequate food and water.

### Dexamethasone gavage in vivo

We prepared a fresh 25 mg/mL stock of water-soluble DEX (Sigma-Aldrich) each day approximately 10 min prior to the time of dosing (i.e., AM and PM). At the time of gavage, DEX was administered based on mouse weight to achieve a dose of 0.5 mg/kg. For gavage, mice were briefly anesthetized with 2% isoflurane and received between 100-200μL solution depending on mouse weight. DEX or vehicle was administered at either ZT4 (morning) or ZT12 (evening) for 6 consecutive days after tumor growth was established at 11 days post-implant. Total dose for mice receiving DEX equaled 3 mg/kg in all treatment groups. Treatment groups are as follows:

– DEX AM: Mice received 0.5mg/kg DEX at ZT4 and vehicle at ZT12 for 6 consecutive days.
– DEX PM: Mice received vehicle at ZT4 and 0.5mg/kg DEX at ZT12 for 6 consecutive days.
– Vehicle: Mice received vehicle at both ZT4 and ZT12 for 6 consecutive days.

### Body weight measurements and survival

To assess tumor burden and disease progression, mice were weighed daily starting on the day of implant. At the end of the experiments, or when mice lost 10% of their body weight from 1^st^ day of tumor implant, these were sacrificed in accordance with IACUC protocols. For tissue collection, Mice were anesthetized with intraperitoneal Avertin (tribromoethanol), then transcardially perfused with 10mL of PBS followed by 10mL of 4% paraformaldehyde (PFA).

### Locomotor activity recording

Mice were singly housed in standard 12 h light/12 h dark, reversed 12h dark/ 12 h light, or constant darkness (DD) conditions in wheel-cages to record locomotor activity in one-minute bins. Wheel-running activity was continuously recorded for 2 weeks before tumor implant to allow for mouse habituation to new cages and single housing, and throughout the end of each experiment.

### Fecal corticosterone analysis

Mice were placed in custom-built collectors, maintained in a temperature-, humidity-, and light-controlled chamber with food and water provided ad libitum to allow for non-invasive measurement of fecal corticosterone from individual mice, a validated method for measuring circadian rhythms in corticosterone release. Mice were allowed to habituate in the cages for at least 7 days before beginning fecal pellet collection. Fecal pellets were collected every hour for a total of 28h. We next removed the collected fecal pellets from the dish, placed them into individual tubes, and baked them at 62 °C until completely dry, typically 2–3 days. Timepoints were binned into 4-hour groups for further processing, and the resultant samples were grounded into a fine powder using a mortar and pestle. We weighed out 20 mg of each sample, added 1 mL of 80% methanol, and agitated for 40 min to extract steroids. After centrifuging, supernatant was transferred to new tubes and placed in a fume hood until methanol fully evaporated (∼7 days). Dried samples were resuspended in 200 µL enzyme-linked immunosorbent assay (ELISA) buffer (Cayman Chemical, Ann Arbor, MI) and then diluted 1:50 in ELISA buffer. We processed the samples in triplicate for corticosterone concentration using the instructions in the ELISA kit (Corticosterone ELISA Kit, Cayman Chemical). Absorbance values were measured using a microplate reader at 415 nm (iMark; BioRad, Hercules, CA) and final corticosterone concentrations (in pg/mL) were determined based on the standard curve and dilution factor.

### Immunohistochemistry

Brains were collected and fixed in 4% PFA overnight at 4°C, then transferred to a sucrose gradient for cryoprotection. Brains were embedded in Tissue-Tek O.C.T. and sectioned in the coronal plane at 40μm using a sliding microtome. Slices were then permeabilized for 30 min with 3% Triton-X in 1x PBS, and blocked for 1 hour with solution containing 10% BSA and 0.3% Triton-X. Primary antibodies were diluted in 2% blocking solution and incubated overnight at 4°C. Sections were then rinsed three times with 1x PBS and incubated in secondary antibody solution in 2% blocking solution for 2 hours at RT. Sections were rinsed 3 times in PBS, mounted with ProLong Gold mounting medium with DAPI (Life Technologies, Carlsbad, CA), and stored in darkness at 4°C until imaging. Microscopy analysis was performed using ImageJ software. All antibodies and reagents used are listed in the Key Resources Table.

## QUANTIFICATION AND STATISTICAL ANALYSIS

### Circadian analyses

The correlation coefficient (CC) of a best-fit circadian cosine function was calculated using ChronoStar 1.0 to assess circadian rhythmicity in GBM cells. CC values above 0.7 were circadian and are reported in figure legends and text. For *in vivo* bioluminescence data, JTK cycle was used to assess circadian rhythmicity, and p value is reported in figure legends and text. A level of p < 0.05 was used to designate circadian rhythmicity.

### Statistics

Data is presented as mean±SEM. Statistical significance of mean differences was determined by either i) Student’s t tests; ii) one-way analysis of variance (one-way ANOVA) with multiple comparisons test; or iii) two-way analysis of variance (two-way ANOVA) with multiple comparisons test. The sample size (n), statistical analysis, and multiple comparisons test used is reported within the figures or legends for each experiment. A level of p < 0.05 was used to designate significant differences. P-values are reported using the following symbolic representation: ns = p > 0.05, ∗ = p ≤ 0.05, ∗∗ = p ≤ 0.01, ∗∗∗ = p ≤ 0.001, ∗∗∗∗ = p ≤ 0.0001. All the statistical analyses were performed using GraphPad Prism (version 10.2.0).

## Supplemental figure titles and legends

**Supplementary Figure 1:**
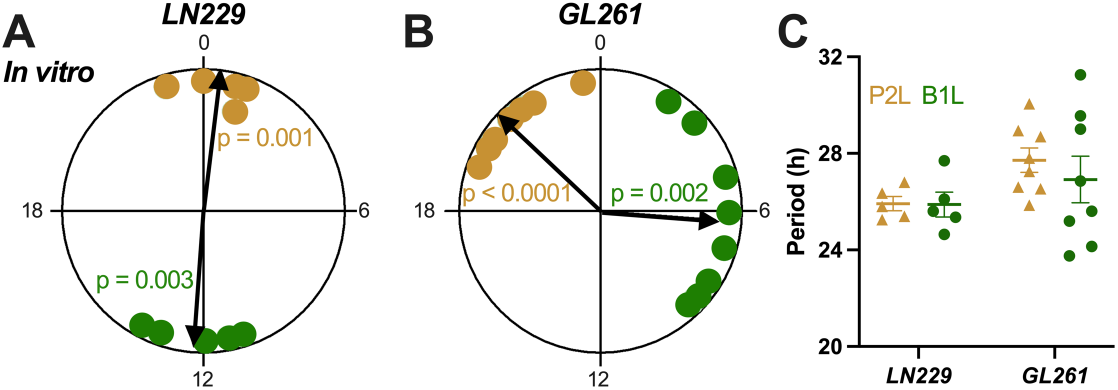
Daily rhythms in *Bmal1* and *Per2* expression peak in anti-phase in GBM cells *in vitro*, Related to Figure 1 (A, B) Rayleigh plots of LN229 and GL261 GBM cells recorded *in vitro* expressing *Bmal1*-luc (green dots; Rayleigh test, p < 0.01) or *Per2*-luc (yellow dots; Rayleigh test, p < 0.01). In this and other Rayleigh plots, arrows point to the mean time of day when the rhythm peaked, and the length of the arrow indicates variability in the data ranging from 0 (peaked at random times) to 1 (all recordings peaked at the same time). (C) Intrinsic period of *Bmal1* and *Per2* expression in LN229 and GL261 GBM cells *in vitro* ranges from 23 to 31 hours, indicating that clock gene expression is circadian.

**Supplementary Figure 2:**
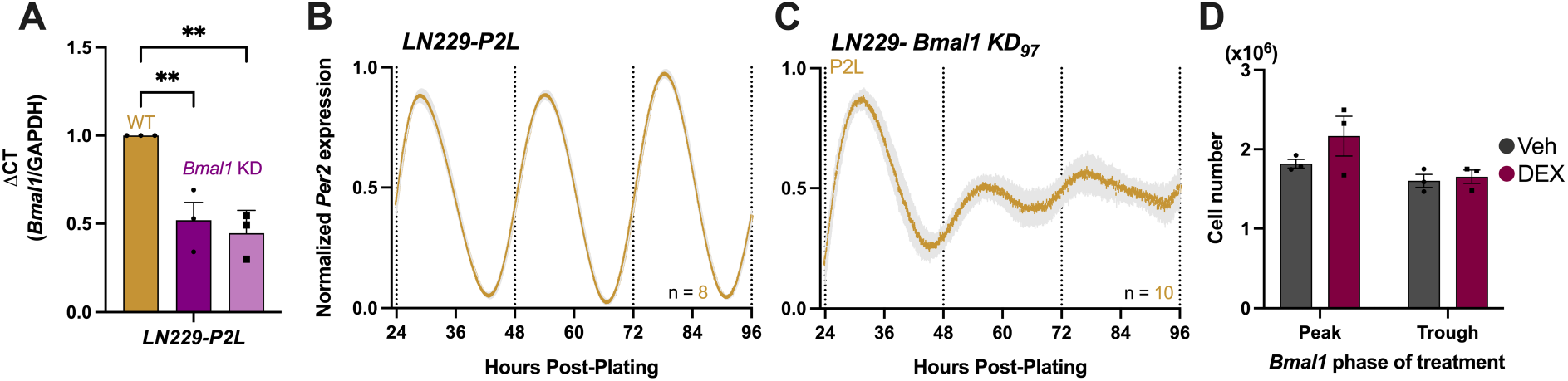
GBM requires an intact circadian clock for daily Dexamethasone-induced tumor growth, Related to Figure 1 (A) shRNA-mediated knockdown reduces *Bmal1* mRNA expression in LN229-P2L GBM cells, using two distinct lentiviral constructs (bright and light purple bars) (mean±SEM, n = 3 per group, one-way ANOVA with Tukey’s multiple comparisons test, **p < 0.01, ns p > 0.05). (B) LN229-P2L cells have intrinsic daily rhythms in *Per2* expression *in vitro* (mean±SEM, all recordings had cosine fits with correlation coefficients, CC > 0.9). (C) An independent *Bmal1* KD lentiviral construct (*Bmal1* KD_97_) reduces the amplitude of *Per2* expression in LN229-P2L cells *in vitro* after 48 hours of recording (mean±SEM, all recordings had cosine fits with correlation coefficients, CC < 0.7). (D) LN229-*Bmal1* KD_97_ cells acutely treated with 100nM Dexamethasone at either the peak or trough of *Bmal1* expression *in vitro* show no differences in cell growth (mean±SEM, n = 3 per group, two-way ANOVA with Šídák’s multiple comparisons test, ns p > 0.05).

**Supplementary Figure 3:**
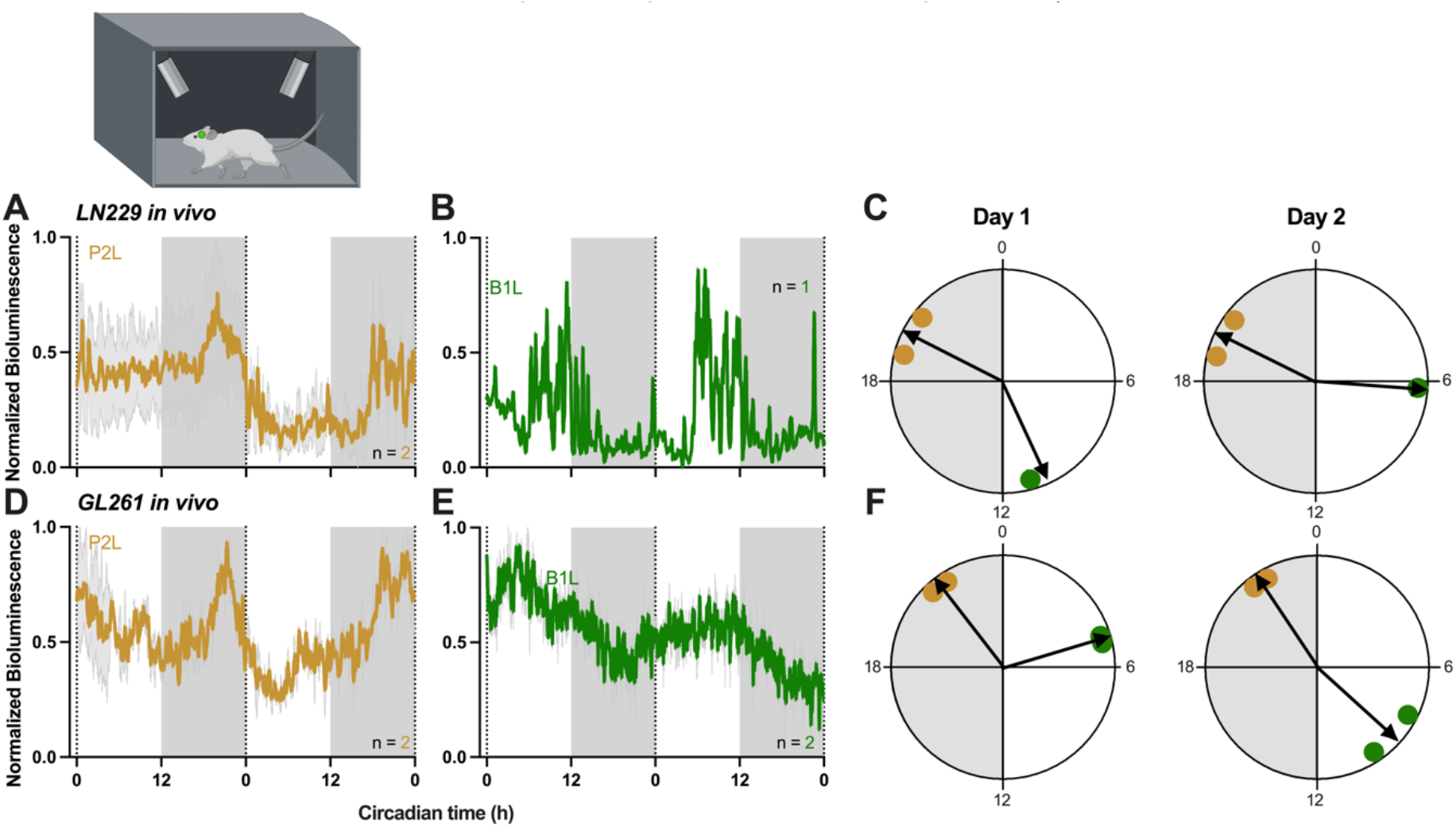
GBM xenografts show circadian rhythms in clock gene expression in freely moving mice, Related to Figure 2 (A-B) Normalized bioluminescence data from freely moving mice housed in DD and implanted with LN229 cells show circadian rhythms in *Per2* (A) and *Bmal1* (B) expression, peaking consistently during the subjective dark or light phase, respectively (mean±SEM, average trace had cosine fit with correlation coefficient, CC > 0.9). (C) Rayleigh plots of LN229 tumors implanted into nude mice, housed in DD, expressing *Bmal1*-luc (green dots; peak time ZT 11 on day 1, ZT 6.5 on day 2, Rayleigh test, p < 0.05) or *Per2*-luc (yellow dots; peak time ZT 20 on day 1 and day 2, Rayleigh test, p < 0.05). Phase was calculated on the first and second day of continuous recording. (D-E) Normalized bioluminescence data from freely moving mice housed in DD and implanted with GL261 cells show circadian rhythms in *Per2* (D) and *Bmal1* (E) expression, peaking consistently during the subjective dark or light phase, respectively (mean±SEM, average trace had cosine fit with correlation coefficient, CC > 0.9). (F) Rayleigh plots of GL261 tumors implanted into C57 mice, housed in DD, expressing *Bmal1*-luc (green dots; peak time ZT 5 on day 1, ZT 9 on day 2, Rayleigh test, p < 0.05) or *Per2*-luc (yellow dots; peak time ZT 21.3 on day 1, ZT 21.5 on day 2, Rayleigh test, p < 0.05). Phase was calculated on the first and second day of continuous recording.

**Supplementary Figure 4:**
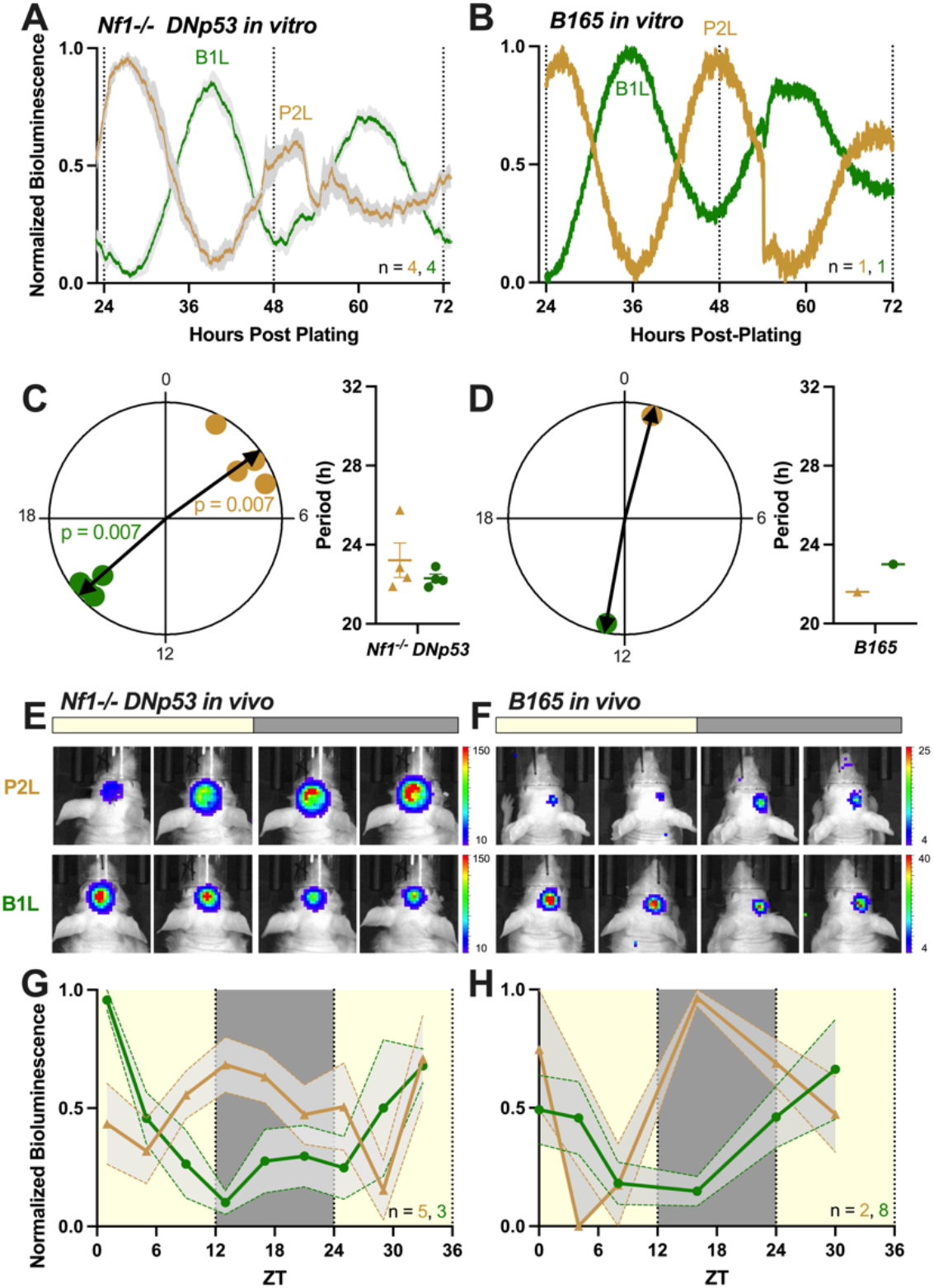
Human GBM cell lines and isolates show intrinsic daily rhythms in *Per2* and *Bmal1* expression independent of tumor origin, Related to Figure 2 (A-B) Human Nf1^-/-^DNp53 and patient-derived B165 GBM cell lines transduced with a *Per2-* or *Bmal1-driven* luciferase reporter (P2L and B1L, respectively), show circadian rhythms in clock gene expression *in vitro* (mean±SEM, all recordings had cosine fits with correlation coefficients, CC > 0.9). (C-D) Rayleigh plots of Nf1^-/-^DNp53 (C) and B165 (D) GBM cells recorded *in vitro* expressing *Bmal1*-luc (green dots; Rayleigh test, p < 0.01) or *Per2*-luc (yellow dots; Rayleigh test, p < 0.01). Clock gene expression peaks in anti-phase in both cell lines. Intrinsic period of *Bmal1* and *Per2* expression in Nf1^-/-^DNp53 (C) and B165 (D) GBM cells *in vitro* ranges from 21 to 25 hours, indicating that clock gene expression is circadian. (E-F) Representative bioluminescence images of tumor xenografts in mice during the day (ZT0-12) and night (ZT12-24) show high *Bmal1* expression during the day, and high *Per2* expression during the night (BLI counts are x10^3^). (G-H) GBM xenografts reliably show peak *Per2* expression at night and *Bmal1* during the day when implanted into nude male or female mice (mean±SEM, average traces scored circadian by JTK cycle p < 0.05).

**Supplementary Figure 5:**
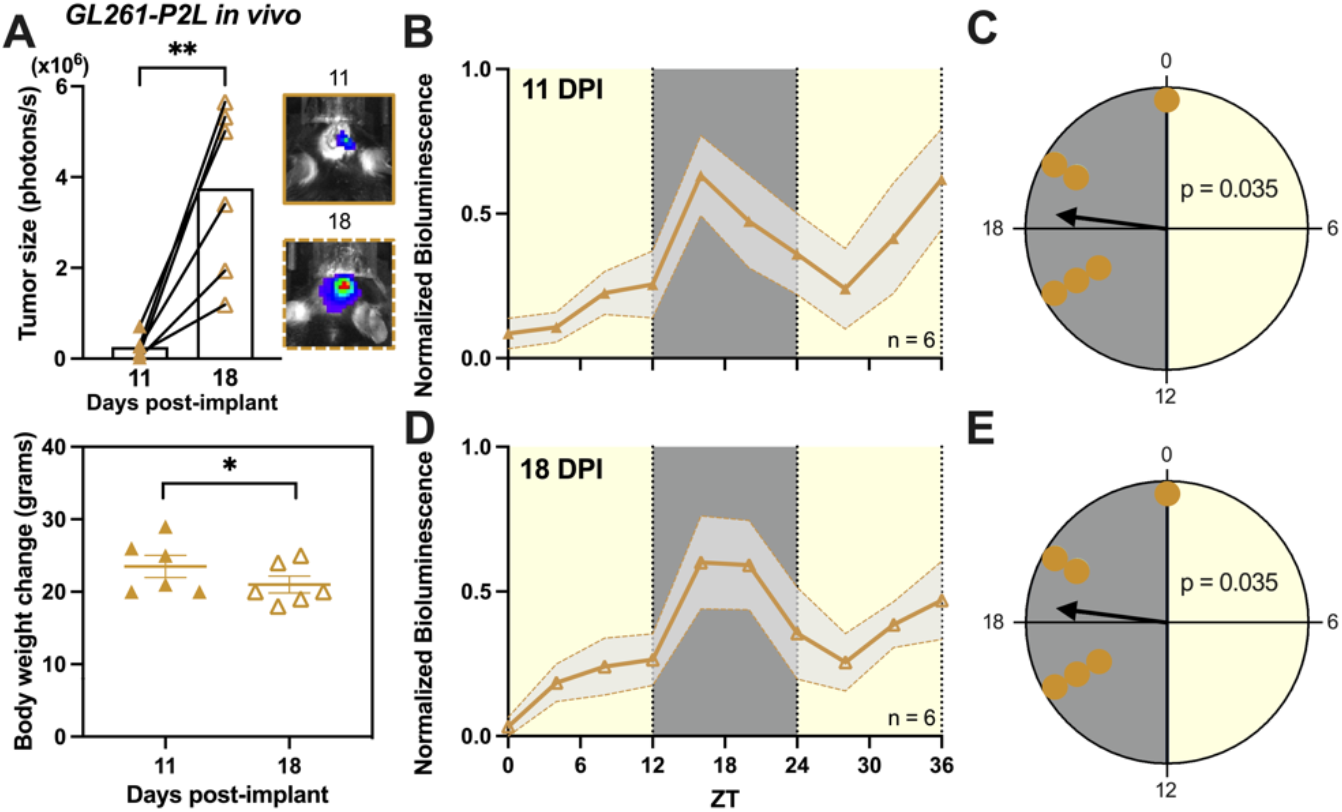
Daily rhythms in *Per2* expression in GBM are sustained throughout tumor progression *in vivo*, Related to Figure 2 (A) Mice bearing GL261-P2L tumors showed significant increase in tumor size (top) and body weight loss (bottom) from 11 to 18 days post-implant (mean±SEM, n = 6, t test, * p < 0.05, ** p < 0.01). (B) GL261-P2L xenografts show peak *Per2* expression at night (ZT16) when imaged at 11 days post-implant (mean±SEM, average traces scored circadian by JTK cycle p < 0.05). (C) Rayleigh plot of GL261-P2L tumor xenografts imaged at 11 days post-implant (yellow dots; peak time ZT 18.5, Rayleigh test, p < 0.05). (D) GL261-P2L xenografts show peak *Per2* expression at night (ZT16) when imaged at 18 days post-implant (mean±SEM, average traces scored circadian by JTK cycle p < 0.05). (E) Rayleigh plot of GL261-P2L tumor xenografts imaged at 18 days post-implant (yellow dots; peak time ZT 18.5, Rayleigh test, p < 0.05).

**Supplementary Figure 6:**
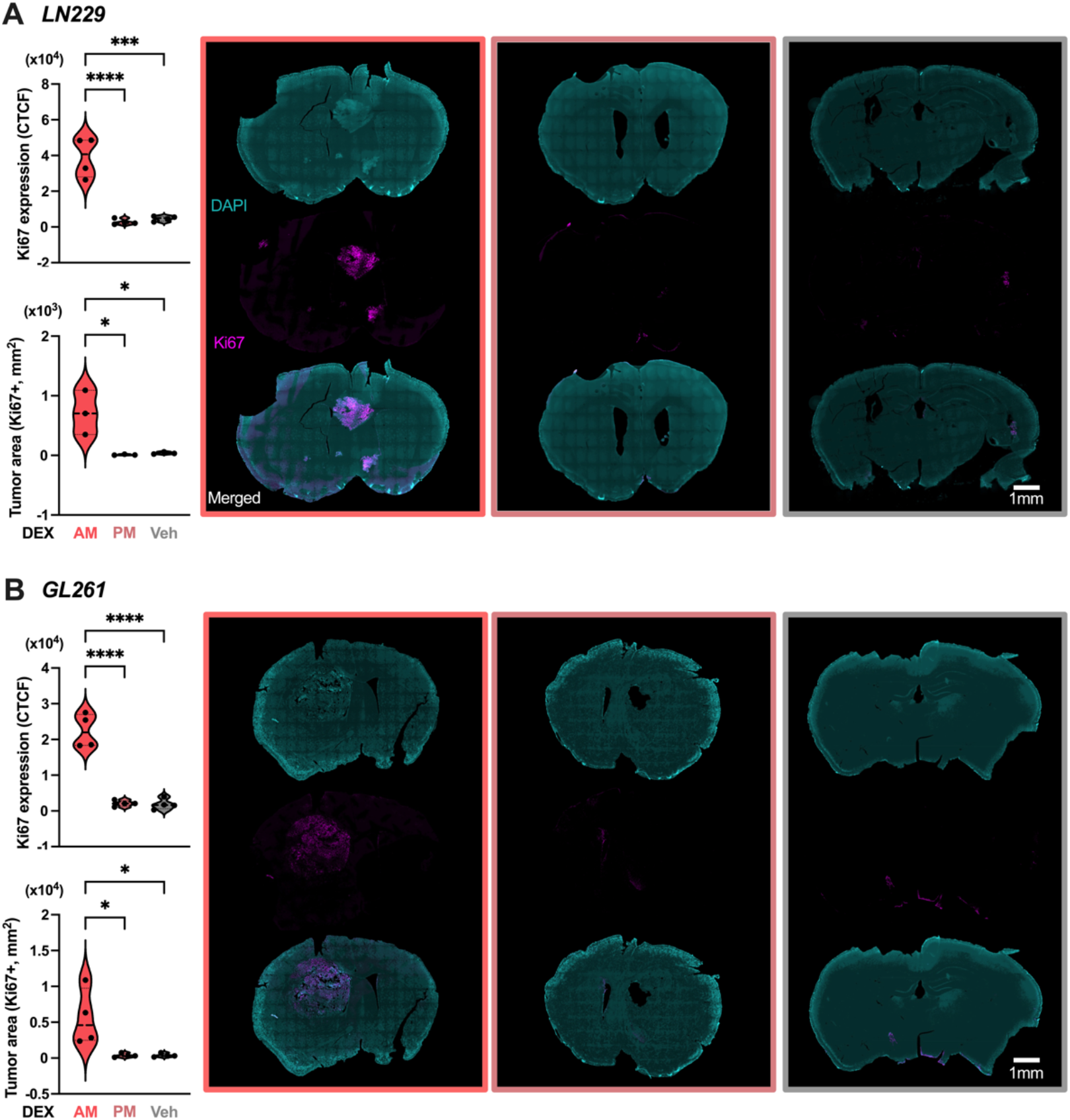
Dexamethasone administration in the morning promotes GBM proliferation and increased tumor area, compared to evening or vehicle treatments *in vivo*, Related to Figure 2 (A-B) Dexamethasone treatment significantly promotes LN229 (A) and GL261 (B) proliferation and increased tumor area *in vivo*, as measured by expression of the proliferation marker Ki67, when administered in the morning (ZT4), compared to evening (ZT12) or vehicle (mean±SEM, scale bar = 1mm, n = 4 per group, one-way ANOVA with Tukey’s multiple comparisons test, *p < 0.05, ***p < 0.001, ****p < 0.0001). DAPI (cyan) staining was used to label nuclei in whole brain sections. Composite images of DAPI and Ki67 (magenta) staining are shown to visualize tumor in the brain.

**Supplementary Figure 7:**
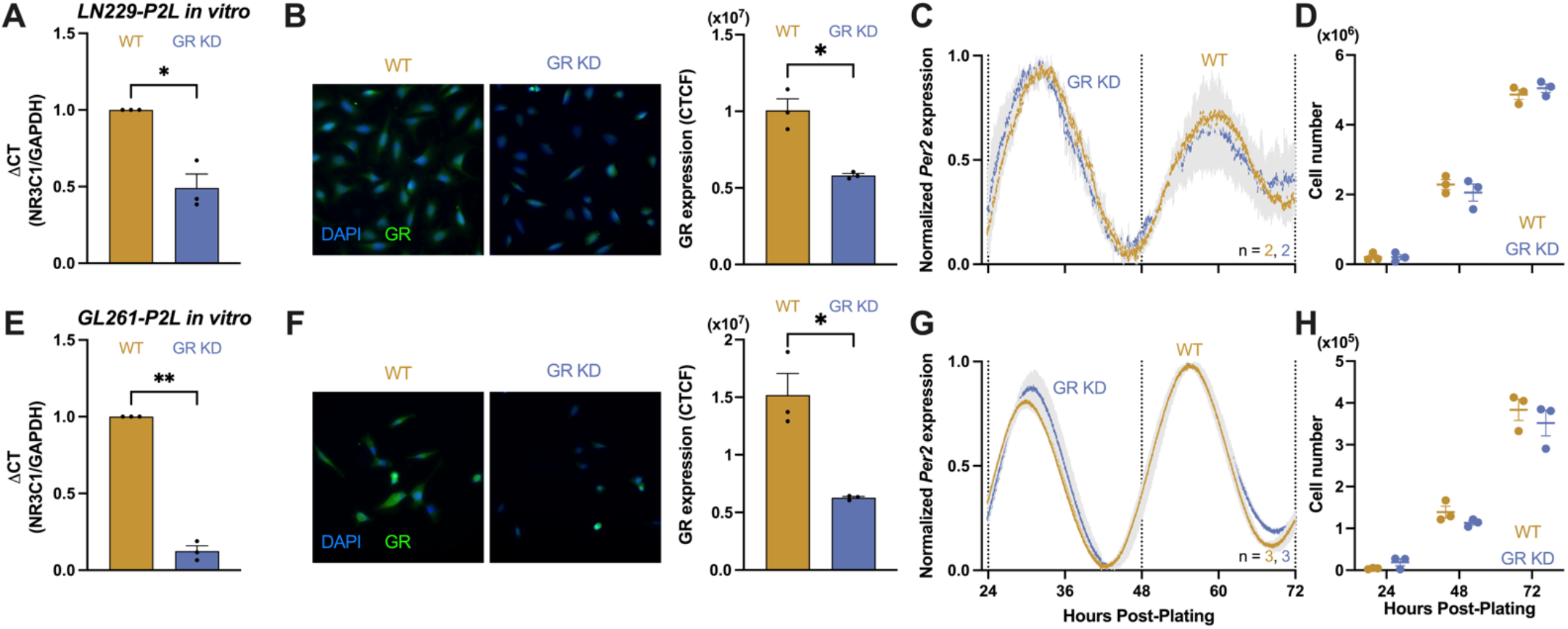
Glucocorticoid receptor knockdown does not affect intrinsic daily rhythms in *Per2* expression or cell growth *in vitro*, Related to Figure 3 (A, E) shRNA-mediated knockdown reduces glucocorticoid receptor (GR) mRNA expression in LN229-P2L and GL261-P2L GBM cells (mean±SEM, n = 3, t test, *p < 0.05, **p < 0.01). (B, F) Representative immunostaining images and quantification of corrected total cell fluorescence (CTCF) intensity shows GBM GR KD cells have significantly decreased GR protein expression, compared to WT (mean±SEM, n = 3, t test, *p < 0.05). DAPI (blue) staining was used to label cell nuclei. Composite images of DAPI and GR (green) staining are shown. (C, G) GR knockdown has no effect on intrinsic daily rhythms in *Per2* expression *in vitro* (mean±SEM, all recordings had cosine fits with correlation coefficients, CC > 0.9). (D, H) GR knockdown has no effect on cell growth over time *in vitro* (mean±SEM, n = 3 per group, two-way ANOVA with Bonferroni’s multiple comparisons test, ns p > 0.05).

**Supplementary Figure 8:**
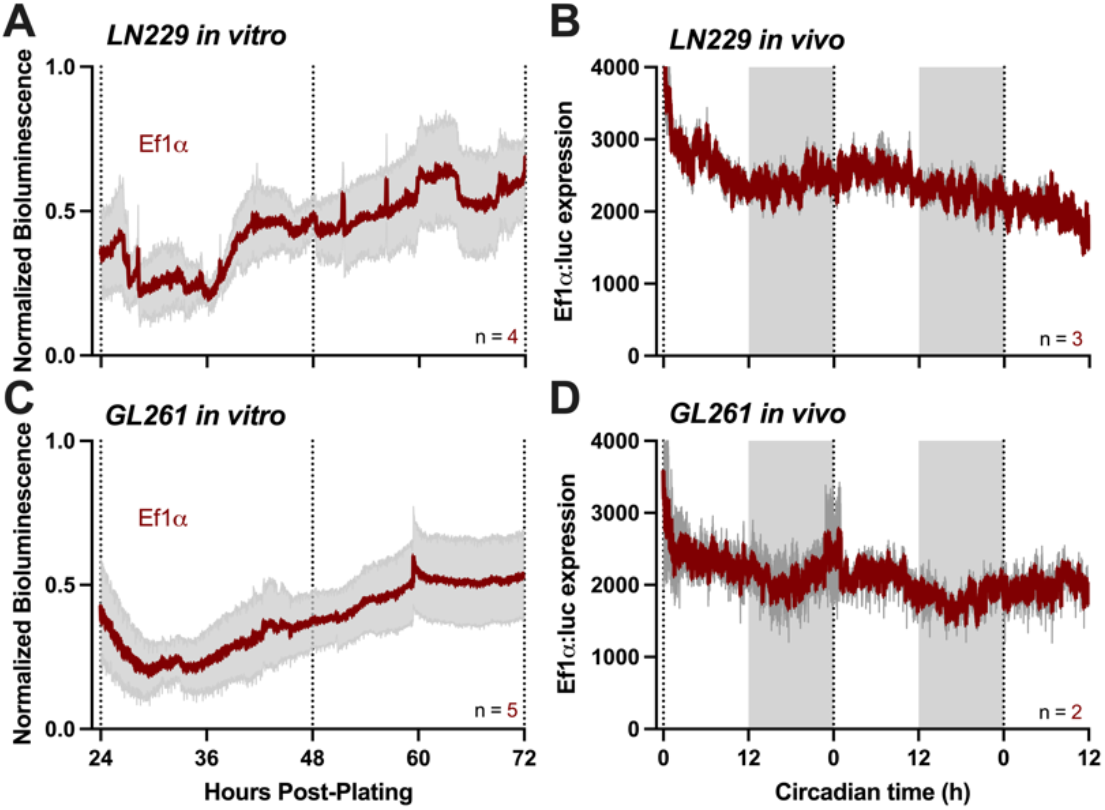
*In vitro* and *in vivo* bioluminescence recordings show no daily rhythms in GBM-Ef1α:luc cells, Related to Figure 3 (A, C) *In vitro* bioluminescence recordings of LN229 and GL261 cells transduced with a *Ef1a*:luc reporter show constitutive *Ef1a* expression over time (mean±SEM, all recordings had cosine fits with correlation coefficients, CC < 0.4). (B, D) Average traces of 60-hour *in vivo* imaging of mice bearing LN229-*Ef1a* and GL261-*Ef1a* GBM tumors two weeks post-implant, in constant darkness, shows constitutive *Ef1a* expression (mean±SEM, all recordings had cosine fits with correlation coefficients, CC < 0.4).

**Supplementary Figure 9:**
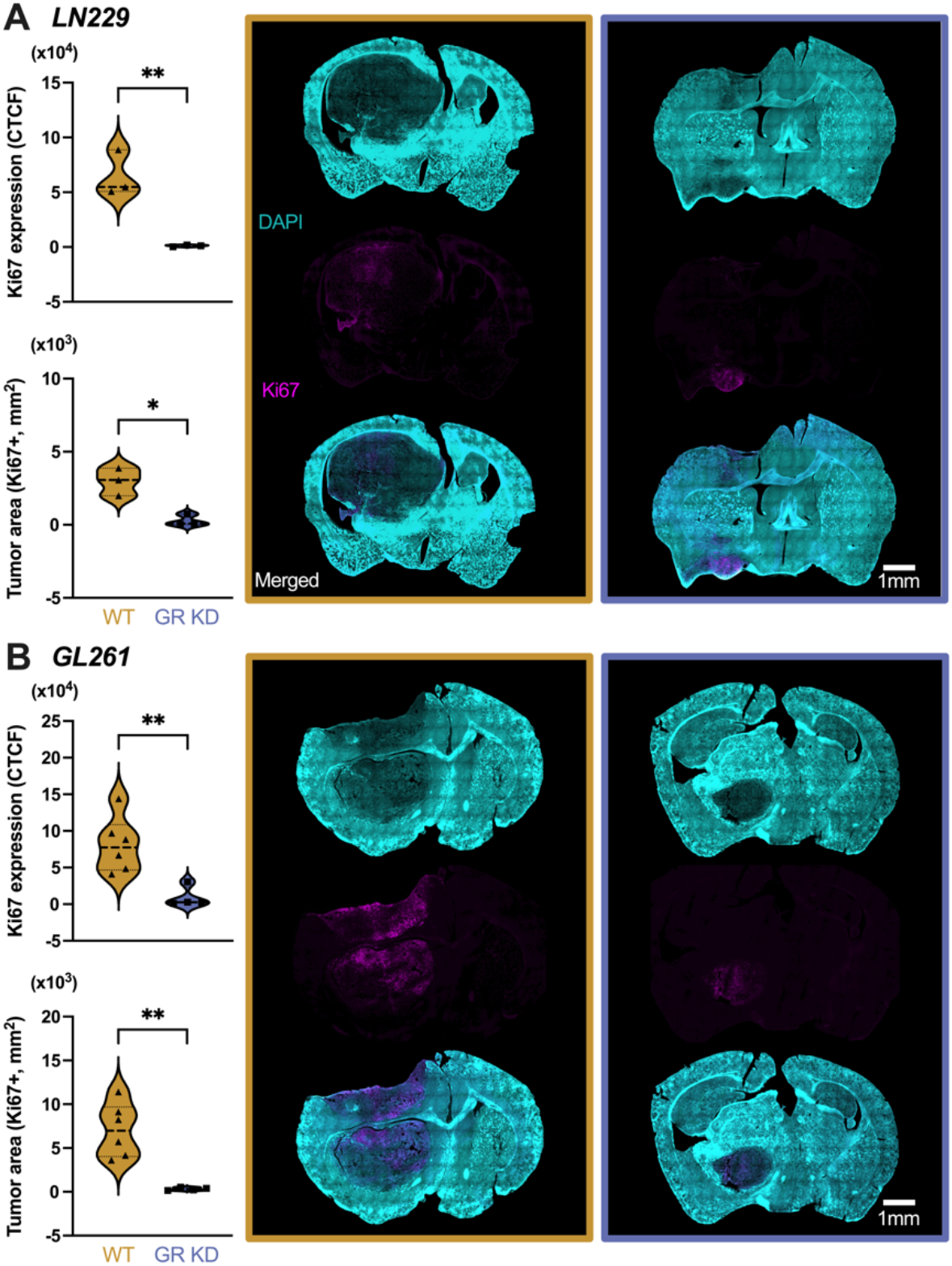
GBM tumors lacking the glucocorticoid receptor show decreased proliferation and smaller tumor area, compared to WT, Related to Figure 3 (A-B) GBM proliferation and tumor area, as measured by Ki67 expression, is higher in mice bearing LN229 (A) and GL261 (B) WT tumors, compared to GR KD (mean±SEM, scale bar = 1mm, n = 3 per group for LN229; n = 6 for GL261, t test, *p < 0.05, **p < 0.01). DAPI (cyan) staining was used to label nuclei in whole brain sections. Composite images of DAPI and Ki67 (magenta) staining are shown to visualize tumor in the brain.

**Supplementary Figure 10:**
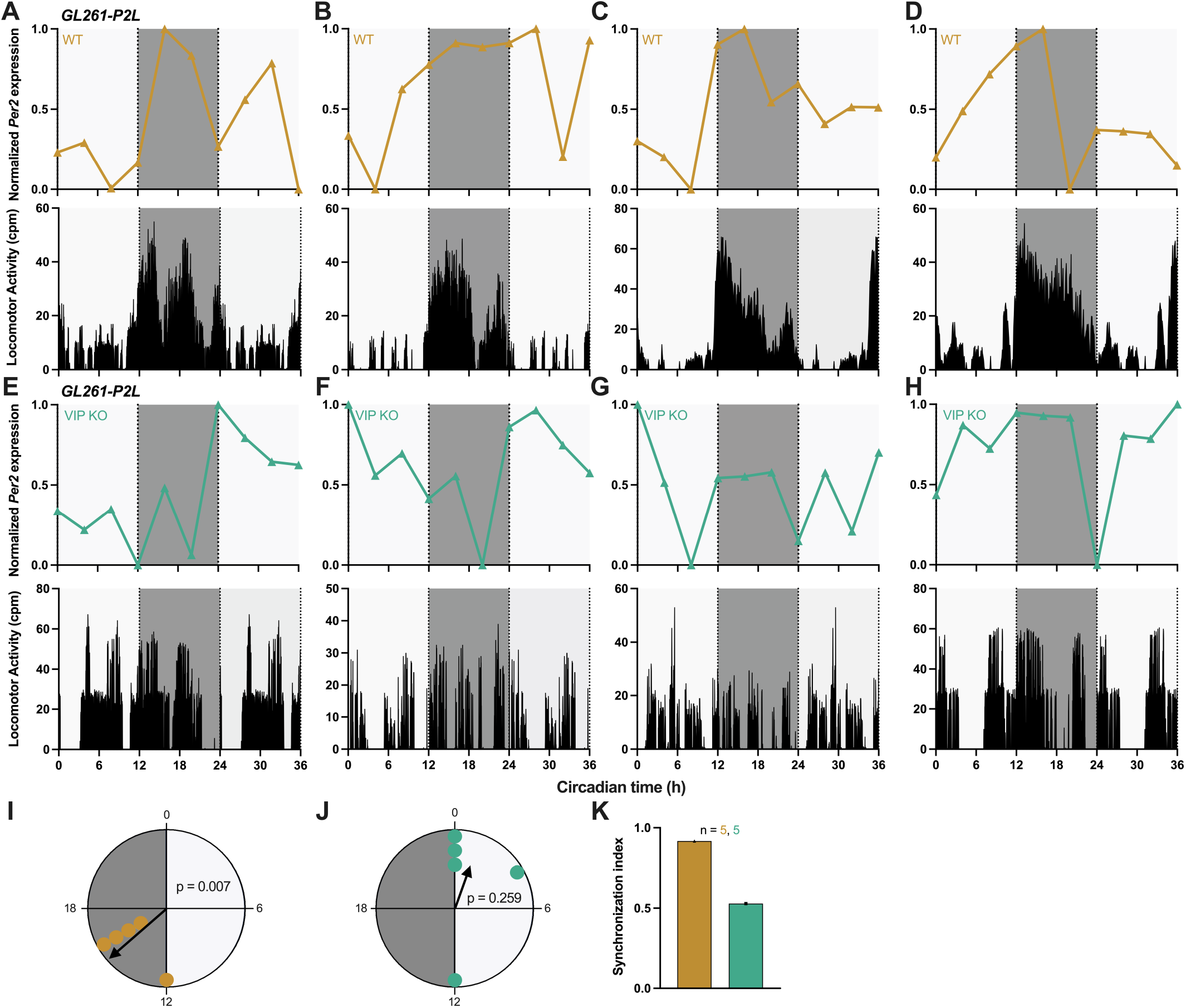
Peak timing of *Per2* synchronizes to host actigraphy in WT, but not arrhythmic VIP KO mice, Related to Figure 6 (A-D) Representative 36-hour *in vivo* bioluminescence imaging (top) and locomotor activity profiles (bottom) of WT mice implanted with GL261-P2L cells, two weeks post-implant and in constant darkness, shows reliable *Per2* peak expression during the subjective dark phase (CT12-24) and rhythmic locomotor activity peaking at CT12 (all traces scored circadian by JTK cycle p < 0.05). (E-H) Representative 36-hour *in vivo* bioluminescence imaging (top) and locomotor activity profiles (bottom) of VIP KO mice implanted with GL261-P2L cells, two weeks post-implant and in constant darkness, shows desynchronized *Per2* peak expression at varying times of day (all traces scored circadian by JTK cycle p < 0.05) and arrhythmic locomotor activity. (I) Rayleigh plot of GL261-P2L tumor xenografts implanted into WT mice shows synchronized *Per2* expression (yellow dots; peak time ZT 15, Rayleigh test, p < 0.05). (J) Rayleigh plot of GL261-P2L tumor xenografts implanted into VIP KO mice shows desynchronized *Per2* expression (green dots; peak time ZT 2, Rayleigh test, p > 0.05). (K) Synchronization index is high in GL261-P2L tumors implanted into WT mice, but decreases in VIP KO mice (mean±SEM).

**Supplementary Figure 11:**
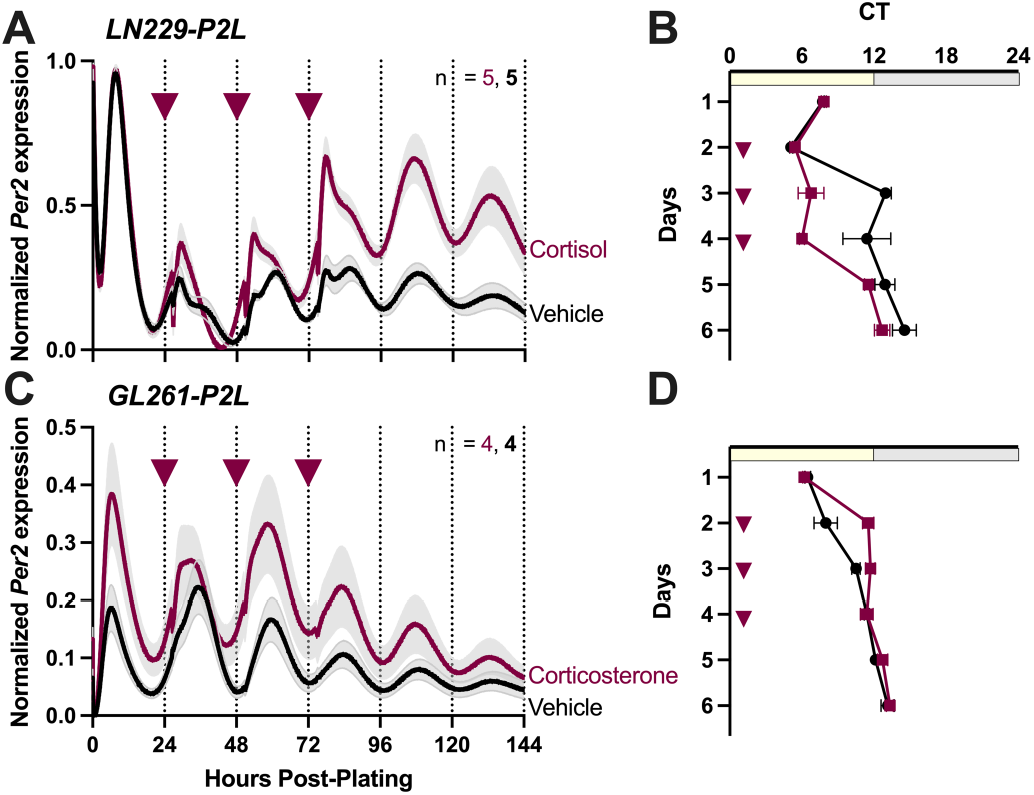
Daily treatment with CORT entrains rhythms in *Per2* expression *in vitro*, Related to Figure 7 (A, C) Representative traces of LN229-P2L and GL261-P2L GBM cells treated chronically for 3 days with Cortisol or Corticosterone, respectively (mean±SEM, all recordings had cosine fits with correlation coefficients, CC > 0.9). (B, D) Chronic daily addition of 10uM CORT induces a stable phase shift in LN229 (6hrs) and GL261 (4hrs), in circadian *Per2* expression, compared to vehicle (mean±SEM).

**Supplementary Figure 12:**
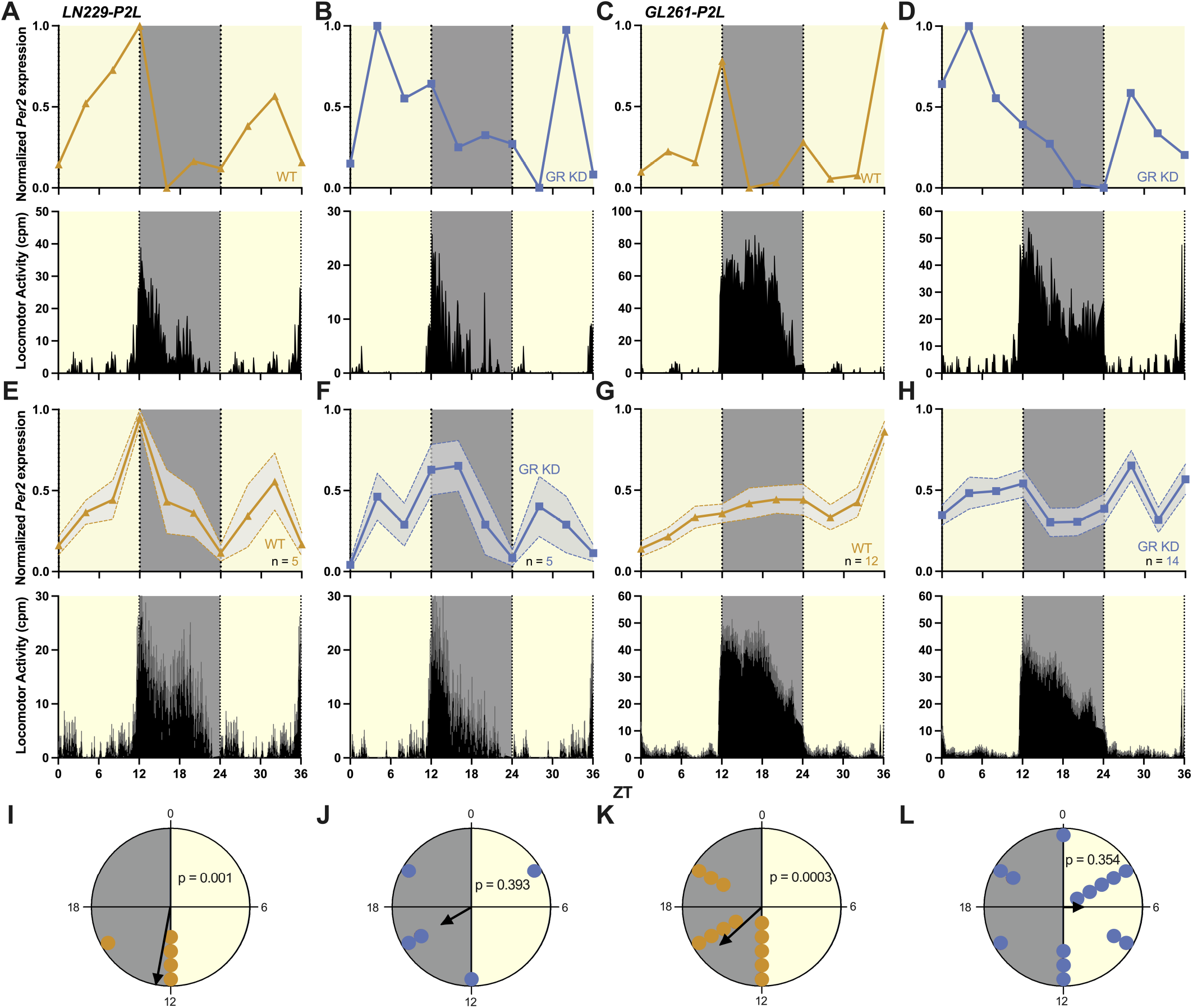
Peak timing of *Per2* synchronizes to host actigraphy in WT, but not GR KD, tumors in a light/dark cycle, Related to Figure 7 (A-D) Representative 36-hour *in vivo* bioluminescence imaging (top) and locomotor activity profiles (bottom) of mice implanted with LN229 WT (A), LN229 GR KD (B), GL261 WT (C), and GL261 GR KD (D) cells, two weeks post-implant and in a standard 12L:12D light schedule. *Per2* expression reliably peaks during the dark phase in WT tumors and in synchrony with locomotor activity onset, but peaks at varying times of day in GR KD tumors (all traces scored circadian by JTK cycle p < 0.05). (E-H) Average traces of 36-hour *in vivo* imaging (top) and locomotor activity profiles (bottom) of mice bearing LN229 WT (I), LN229 GR KD (J), GL261 WT (K), and GL261 GR KD (L) tumors two weeks post-implant. *Per2* expression peaks during the dark phase and in synchrony with locomotor activity onset in WT, but at varying times of day in GR KD tumors (mean±SEM). (I, K) Rayleigh plots of *Per2* expression in LN229 (I) and GL261 (K) WT tumors from tumor-bearing mice housed in LD show reliable peak expression during the dark phase (yellow dots represent a single mouse; peak time ZT 13.06 and 15.38 for LN229 and GL261, respectively; Rayleigh test, p < 0.001). (J, L) Rayleigh plots of *Per2* expression in LN229 (J) and GL261 (L) GR KD tumors from tumor-bearing mice housed in LD show desynchronized peak expression (blue dots represent a single mouse; peak time couldn’t be calculated by Rayleigh statistics; Rayleigh test, p > 0.05).

